# Natural variation in an NLR pair confers thermostable resistance to a devastating bacterial pathogen

**DOI:** 10.64898/2025.12.15.694456

**Authors:** Nathalie Aoun, Mandeep Singh, Chacko Jobichen, Natsumi Maruta, Caitlin Griffiths, Elena Dangla, Heloise Demont, Sébastien Carrere, Jérôme Gouzy, Rémi Duflos, Matilda Zaffuto, Marta Marchetti, Angela M. Hancock, Maud Bernoux, Bostjan Kobe, Laurent Deslandes, Fabrice Roux, Richard Berthomé

**Affiliations:** Department of Plant Pathology, University of California Davis, Davis, CA, USA; Univ Toulouse, INRAE, CNRS, LIPME, Castanet-Tolosan, France; School of Chemistry and Molecular Biosciences, Institute for Molecular Bioscience and Australian Infectious Diseases Research Centre, The University of Queensland, Brisbane, QLD, Australia; Centre for Bacterial Cell Biology, Biosciences Institute, Faculty of Medical Sciences, Newcastle University, Newcastle upon Tyne, United Kingdom; Department of Botany and Plant Pathology, Purdue University, West Lafayette 47907 IN, USA

## Abstract

Climate change reshapes host-pathogen interactions by increasing pathogen aggressiveness and weakening plant immune responses, particularly under heat stress. Nucleotide-binding leucine-rich repeat (NLR) immune receptors, key players in pathogen recognition, are altered under elevated temperatures in both plants and mammals, posing a major challenge to disease resistance. In *Arabidopsis thaliana,* the well-described RPS4/RRS1 NLR pair confers resistance to the worldwide devastating phytopathogenic *Ralstonia pseudosolanacearum*. Here, by combining the exploration of natural genetic variation with genetic mapping, polymorphism analysis, structural modeling, and functional complementation, we demonstrated that the paucity of full thermostable resistance is primarily associated with a unique *RPS4/RRS1* haplotype carrying key substitutions in leucine-rich repeat domains. These findings reveal natural genetic diversity as a source of thermostable resistance, highlighting promising opportunities to engineer climate-resilient plants.

## Main text

Climatic fluctuations linked to anthropogenic activities have a profound impact on the functioning of ecosystems and the geographical distribution of pathogens, which together increase the worldwide frequency and severity of epidemics affecting humans, animals, and plants (*1*). In crops, such epidemics result in devastating economic losses and put our global food security at risk (*2*). Among the climatic parameters involved, global average temperatures are expected to reach or, likely, surpass the 1.5°C threshold for the 2021-2040 period, according to IPCC (Intergovernmental Panel on Climate Change) (*1*). Such temperature elevation has been demonstrated to deeply impact organismal immunity through the impairment of key resistance mechanisms, particularly those involving plant nucleotide-binding leucine-rich repeat (NLR) immune receptors (*3*, *4*). To mitigate these global disease challenges, it is crucial to characterize the genetic mechanisms underlying the natural genetic variation of plant defense responses to pathogens at elevated temperatures, to identify full thermostable resistances.

To address this goal, we investigated under heat stress conditions the interaction between *Ralstonia pseudosolanacearum,* one of the most devastating plant pathogenic bacteria worldwide (*5*), and the model plant *Arabidopsis thaliana*. *Ralstonia solanacearum* species complex (RSSC) are root-invading pathogens causing bacterial wilt disease on a broad host range, including over 500 crops and wild species across 87 botanical families (*6*). In *A. thaliana*, RESISTANCE TO RALSTONIA SOLANACEARUM1 (RRS1) and RESISTANCE TO PSEUDOMONAS SYRINGAE4 (RPS4) are a pair of Toll/interleukin-1 receptor-nucleotide-binding site leucine-rich repeat (TIR-NB-LRR, TNL) NLR immune receptors, in which RRS1 functions as a pathogen sensor as wells as abiotic signals related to humidity (*7*), while RPS4 as a helper (or executor) for activation of immune responses. In this pair, RRS1 and RPS4 form a heterodimeric complex that cooperatively recognizes multiple effector proteins (*8–11*), including the YopJ family acetyltransferase PopP2 from *R. pseudosolanacearum* (*12*, *13*). PopP2 acetylation of the C-terminal WRKY domain of RRS1 that mimics the effector’s true targets causes derepression of the NLR complex and activation of the RPS4-dependent immunity (*12–14*).

Previously, the phenotyping of a worldwide *A. thaliana* genome-wide association (GWA) mapping panel in response to root inoculation with the *R. pseudosolanacearum* GMI1000 reference strain was performed at permissive (27°C) and elevated (30°C) temperatures to investigate the impact of temperature elevation on disease response (*15*). Almost all tested natural accessions became more susceptible 10 days after inoculation (dai) at 30°C, compared to 27°C. While the major resistance QTL (quantitative trait locus) identified at 27°C corresponds to the *RPS4/RRS1* locus (*15*), an increase in temperature to 30°C revealed a highly polygenic architecture with no QTL located on the *RPS4/RRS1* pair, indicating that RPS4/RRS1-mediated immunity was negatively affected at elevated temperatures. Functional validation of minor QTLs uncovered unexpected players conferring resistance at 30°C, including *AtSSL4* and *AtSDS,* which encode a strictosidine synthase-like protein and an atypical meiotic cyclin, respectively (*15*, *16*). It is striking that neither of these genes had been previously linked to plant immunity. Notably, within the GWA mapping panel, the Eden-1 accession (sampled from Sweden) was the only one that remained symptom-free at 30°C when inoculated with GMI1000.

To identify additional accessions with such a full thermostable disease resistance to *R. pseudosolanacearum* and characterize the underlying genetic bases, we investigated the natural genetic variation in disease symptoms caused by GMI1000 at 27°C and 30°C in an additional panel of 103 worldwide accessions. Many accessions of this panel originate from geographic regions underrepresented in our previous screening panel (e.g., Africa, Central Asia, Iberian Peninsula, and the Middle East) (Fig. 1A, Table S1). While most accessions were more susceptible to GMI1000 at 30°C compared to 27°C, the variability of disease response at this temperature was greater than in our previous worldwide panel (coefficient of variation of the new panel = 0.277, coefficient of variation of the previous panel = 0.114) (Fig. 1B). Among the 103 accessions tested, Anz-0 (sampled from Iran) was the only accession phenotypically similar to Eden-1, with no visible disease wilting symptoms observed at 30°C at 10 dai (Fig. 1A-B, Table S2).

**Fig. 1.**
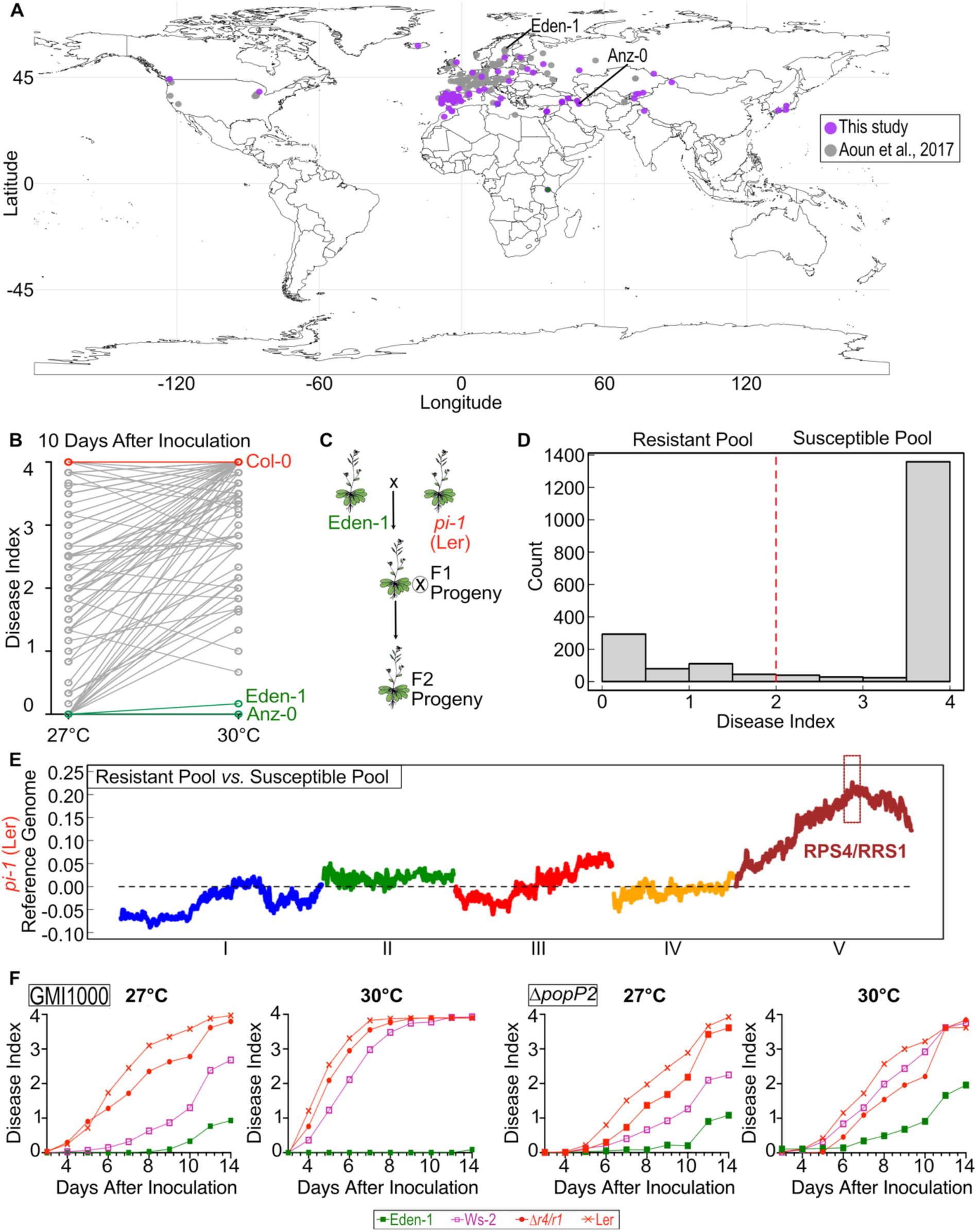
Natural variation in *Arabidopsis thaliana* reveals thermostable disease resistance to *Ralstonia pseudosolanacearum*. (**A**) Geographic origin of *A. thaliana* accessions screened for response to GMI1000 at 27°C and 30°C in previous (Aoun et al., 2017, grey) and current studies (purple). (**B**) Within-group adjusted mean disease index of accessions evaluated in this study, shows natural variation in response to GMI1000 across temperatures. Eden-1 and Anz-0 exhibited thermostable disease resistance at 30°C. (**C**) Crossing of the resistant Eden-1 and susceptible *pi-1* mutant (Ler-0 background) to generate the F2 segregated population in the bulk segregant analysis approach. (**D**) Distribution of disease index in F2 plants between resistant and susceptible pools in response to GMI1000 infection at 30°C. **(E)** Allele frequency differences between resistant and susceptible pools mapped on the *pi-1* genome revealed a major QTL at the *RPS4/RRS1* locus. (**F**) Disease assays with GMI1000 and Δ*popP2* knockout mutant strains on Eden-1, Ler-0, Ws-2, and the *rps4-21/rrs1-1* double mutant show that Eden-1 thermostable disease resistance is *PopP2*-dependent.

We first characterized the genetic architecture of the thermostable disease resistance of Eden-1 by adopting a bulk segregant analysis (BSA) approach. Eden-1 was crossed with the susceptible *PISTILLATA-1* (*pi-1*) male sterile EMS (ethyl methanesulfonate) mutant in the Landsberg erecta (Ler) genetic background (Fig. 1C). The resulting F2 segregating population (N=1978 plants) was root-inoculated with GMI1000 at 30°C. At 10 dai, 73.3% and 26.7% of the F2 plants were susceptible and resistant to GMI1000, respectively (Fig. 1D). These relative percentages did not deviate significantly from the ‘1:3 (resistant:susceptible)’ segregating ratio (χ2 = 3.03 < 3.84; α = 5%), suggesting that Eden-1’s thermostable resistance was mainly governed by a single recessive nuclear locus. To map this major locus, we adopted a mapping-by-sequencing approach by sequencing one pool of F2 resistant plants (N = 167) and seven pools of F2 susceptible plants (N=7 x 167) (Supplementary methods). We mapped the corresponding Illumina reads sequencing data on the *de novo* and annotated PacBio long-read-based reference genomes of Eden-1 and *pi-1*. Among the eight pools considered, we identified 354,763 and 316,504 SNPs on Eden-1 and *pi-1* reference genomes, respectively (Supplementary methods). The differentiation in allele frequency between the resistant pool and the seven susceptible pools revealed a major QTL located at the end of chromosome V, along with four minor QTLs located on chromosomes I and III (Figs. 1E, S1). In the major QTL region, the top SNP was located (i) within the *RPS4/RRS1* pair when the reads were mapped on the *pi-1* genome, and (ii) 62 kb downstream of the *RPS4/RRS1* pair when the reads were mapped on the Eden-1 genome (Supplementary methods).

To assess the role of *RPS4/RRS1* in Eden-1 thermostable resistance, we conducted inoculation assays at 27°C and 30°C with the GMI1000 strain and a Δ*popP2* knockout mutant strain that is unable to trigger activation of the RPS4/RRS1-mediated immunity (*10*, *12*). We inoculated Eden-1, the Ler accession susceptible at both temperatures, the Ws-2 accession resistant at 27°C but susceptible at 30°C, and the *rps4-21/rrs1-1* double knockout mutant, originating from the Ws-2 background (*17*). As additional controls, the Col-0 accession susceptible at both temperatures and the Nd-1 accession resistant at 27°C but susceptible at 30°C were included in these assays (Fig. S2, Table S3). In contrast to Ws-2 and Nd-1 whose *RRS1* alleles confer recognition of PopP2, PopP2 is not recognized by RRS1 in Col-0, due to the truncation of the C-terminal extension beyond the WRKY domain in RRS1 (*18*, *19*). Due to their inability to recognize PopP2, *rps4-21/rrs1-1* and Col-0 plants became fully wilted in response to the inoculated strains at both temperatures (Fig. 1F, Tables S3-S4). In response to GMI1000, while the Ws-2, Nd-1 and Eden-1 plants were fully resistant at 27°C, only Eden-1 plants remained symptom-free at 30°C (Fig. 1F, Fig. S2). In all cases, the observed resistance response was found to depend on PopP2 (Fig. 1F). Collectively, these data indicate that, while the RPS4/RRS1-mediated immunity is thermolabile in Ws-2 and Nd-1, the RPS4/RRS1 pair in Eden-1 (hereinafter designated as RPS4/RRS1^Eden-1^) confers PopP2 recognition and allows this accession to mount a robust immune response that remains efficient at 30°C (Fig. 1F).

To identify the genetic polymorphisms responsible for RPS4/RRS1^Eden-1^ thermostability, we analyzed long-read-based sequences of the *RPS4/RRS1* locus across 114 Arabidopsis natural accessions, which were scored for disease symptoms in response to GMI1000 at 27°C and 30°C (Table S2). The phylogenetic tree based on 348 polymorphisms identified in the sequences of the *RPS4/RRS1* genomic region (86 in *RPS4,* 18 in the intergenic region, and 244 in *RRS1*) (Tables S5-S7, Fig. 2A) does not align with either the geographic origin of the accessions or the impact of temperature elevation on disease response (Fig. 2B). Six polymorphisms were specific to Eden-1: (i) two synonymous polymorphisms and one nonsynonymous polymorphism in *RPS4*, (ii) one in the intergenic region, and (iii) one synonymous polymorphism and one nonsynonymous polymorphism in *RRS1* (Tables S5-S7). The nonsynonymous polymorphism in *RPS4* is located in exon 4, which encodes the beginning of the leucine-rich repeat (LRR) domain; it corresponds to a T-to-C transition, resulting in a cysteine-to-arginine substitution at position 674 (C674R) in RPS4^Eden-1^ protein (Fig. 2C, Tables S5). The nonsynonymous polymorphism in *RRS1* is located in exon 3, which encodes the beginning of the LRR domain; it corresponds to a G-to-A transition, resulting in an alanine-to-threonine substitution at position 641 (A641T) in the RRS1^Eden-1^ protein (Fig. 2C, Table S7). Interestingly, in addition to the well-established role of the LRR domain in mediating pathogen recognition specificity (*20*, *21*), our results suggest that it may also contribute to modulating plant responses to multiple stress conditions. These findings raised the intriguing possibility that thermostable resistance may arise from accession-specific polymorphisms at the *RPS4/RRS1* sequence. Notably, the naturally occurring A641T non-synonymous polymorphism is located in a genomic region similar to that of an EMS-induced point mutation reported in the *int102/snc1-1* mutant of *A. thaliana*. This substitution corresponds to a glutamic acid-to-lysine change at position 640 (E640K) in the LRR domain of the autoactive SNC1 protein, converting this thermolabile NLR into a thermostable one. This point mutation was indeed demonstrated to provide temperature-resilient immunity mediated by autoactive *snc1-1* (*22*). A similar mutation introduced into the tobacco NLR gene *N,* resulting in a tyrosine-to-lysine substitution at position 646 (Y646K), was also found to confer heat-stable defense responses upon recognition of the p50 elicitor from tobacco mosaic virus (*23*).

**Fig. 2.**
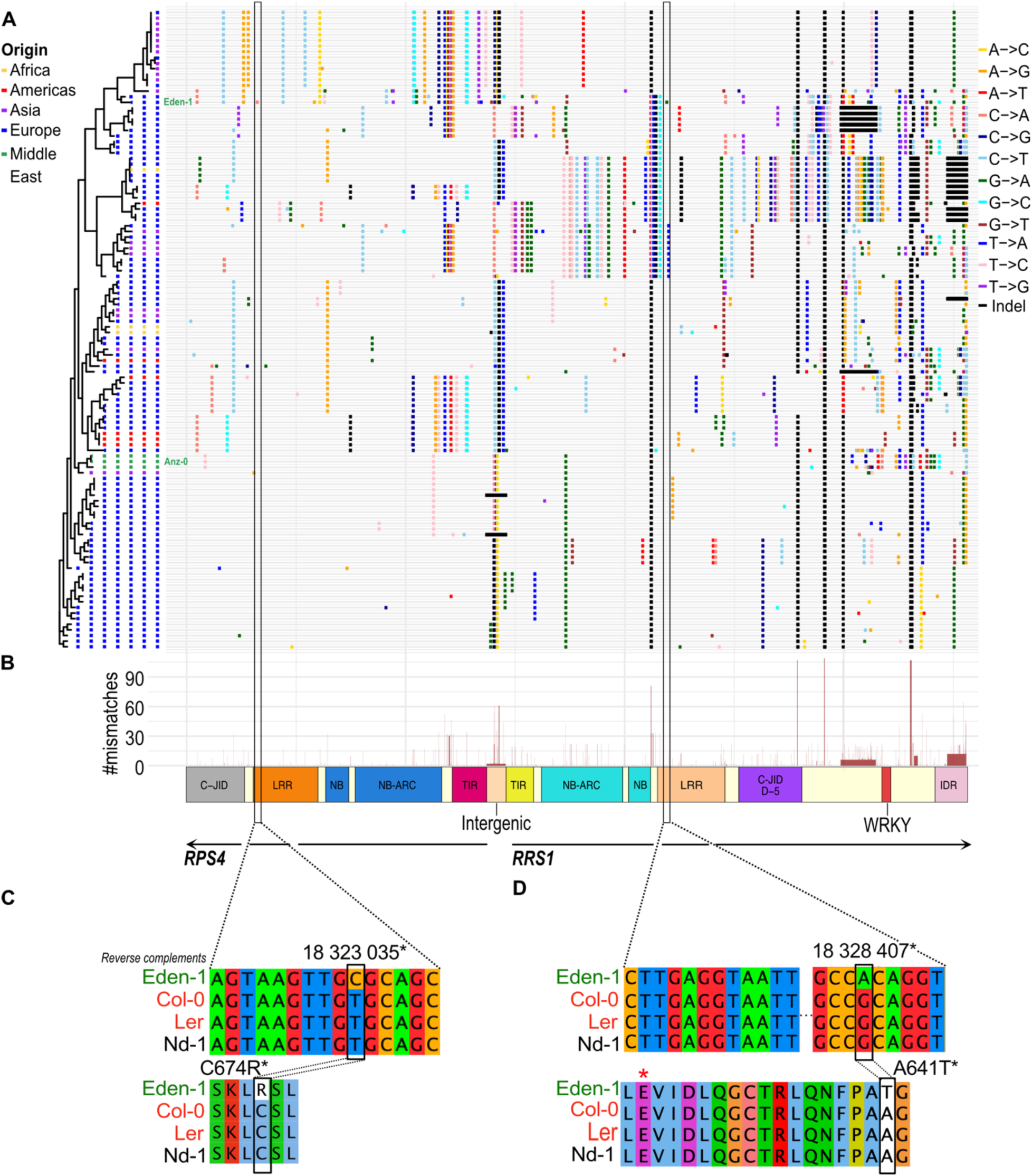
Eden-1 has unique polymorphisms in the LRR domains of *RPS4* and *RRS1* NLR resistance genes. (**A**) Phylogenetic tree of *A. thaliana* accessions constructed from nucleotide sequences of *RPS4*, *RRS1*, and their intergenic region. Branch colors indicate geographic origin: Africa, Americas, Asia, Europe, and the Middle East. The adjacent heatmap displays nucleotide substitutions and indels across accessions. The vertical rectangles indicate the location of Eden-1’s unique polymorphisms in the nucleotide sequences of *RPS4* and *RRS1* encoding their LRR domains. (**B**) Counts of nucleotide mismatches across *RPS4/RRS1*. (**C**) and (**D**) Detailed view of polymorphic sites across Eden-1, Col-0, Ler, and Nd-1 (nucleotide and protein sequences). Non-synonymous mutations (**C**) C674R in RPS4 and (**D**) A641T in RRS1 are highlighted. Black asterisks mark genomic positions in Eden-1. Numbers represent the genomic location of the DNA sequence. The red asterisk marks the residue corresponding to E640 in SNC1; substitution of this residue to lysine (E640K) in the *int102/snc1-1* mutant confers thermostability to SNC1.

To investigate which of Eden-1’s amino-acid changes might confer thermostable resistance, we turned to structure prediction of the RPS4/RRS1 complex. As no experimental three-dimensional structure is available for the RPS4/RRS1 complex, we employed structure prediction using AlphaFold 3 (*24*) to explore potential molecular mechanisms responsible for thermostable resistance. Singleton NLRs confer resistance by converting from a presumably monomeric inactive state, upon pathogen effector recognition, to an active tetrameric resistosome complex (*25*). While no full-length, high-resolution structures of intact paired NLR complexes exist, domain-level structural and related interaction data can be exploited. For RPS4/RRS1, effector recognition has been proposed to drive a transition from a heterodimer to a dimer-of-heterodimers stoichiometry (*14*, *26*, *27*), although an alternative dimer-of-heterodimers configuration without an apparent change in size has more recently been suggested (*28*). The polymorphisms responsible for thermostable resistance could affect both the inactive and resistosome states. Due to limitations of current prediction tools, we were only able to model the heterodimer configuration and assess the effects of temperature elevation on the stabilization of the inactive state of the complex.

Remarkably, the predicted structures of the RPS4/RRS1 complex of the thermolabile (Nd-1) and thermostable (Eden-1) accessions consistently showed distinct conformations that may underpin the corresponding functional differences (Fig. 3). Both proteins contain TIR, NB-ARC (nucleotide-binding adaptor shared by APAF-1, R proteins, and CED-4), LRR, and C-JID (C-terminal jelly-roll/Ig-like domain) domains, with RRS1 containing an additional undefined semi-structured region (D-5) adjacent to C-JID, followed by WRKY and IDR (intrinsically disordered region) domains at the C-terminus (Fig. 2B, S3). Within the predicted complex, the TIR domains showed decent concordance with the experimentally resolved PDB structure (ID: 4C6T) of the *A. thaliana* RPS4/RRS1 TIR assembly (Fig. S3). For Nd-1, the modelled structural ensembles mainly showed the LRR and adjacent C-terminal domain regions of both RRS1 and RPS4 positioned away from their corresponding N-terminal TIR domains (Fig. 3A), constituting an “open” conformational state. By contrast, within the Eden-1 structural ensembles, RRS1^Eden-1^ consistently showed the LRR and adjacent D-5 domain regions reoriented towards the N-terminal TIR regions of the complex, with its C-terminal WRKY domain positioned in direct vicinity of the RRS1 TIR domain, constituting a “closed” conformational state (Fig. 3B). This closed, contact-rich structural reorientation plausibly enhances the thermostability profile of Eden-1 by tightening conformational entropy and distributing thermal fluctuations across a denser network of non-covalent contacts, potentially stabilizing the complex and providing one of the possible explanations for the thermostability. Moreover, these compact Eden-1 ensembles may bury solvent-exposed regions and thereby resist heat-induced unfolding, potentially shifting the overall functional equilibria of the system. Such changes could influence active complex assembly, alter subcellular localization, and/or modulate targeting during pathogen attack (*29–32*). These mechanistic links and conformational differences are supported herein by calculations of their energy scores using MutationExplorer (*33*) (Figs. 3, S4, Table S8), indicating that the reorganized Eden-1 complex is a lower-energy, more stable conformation.

**Fig. 3.**
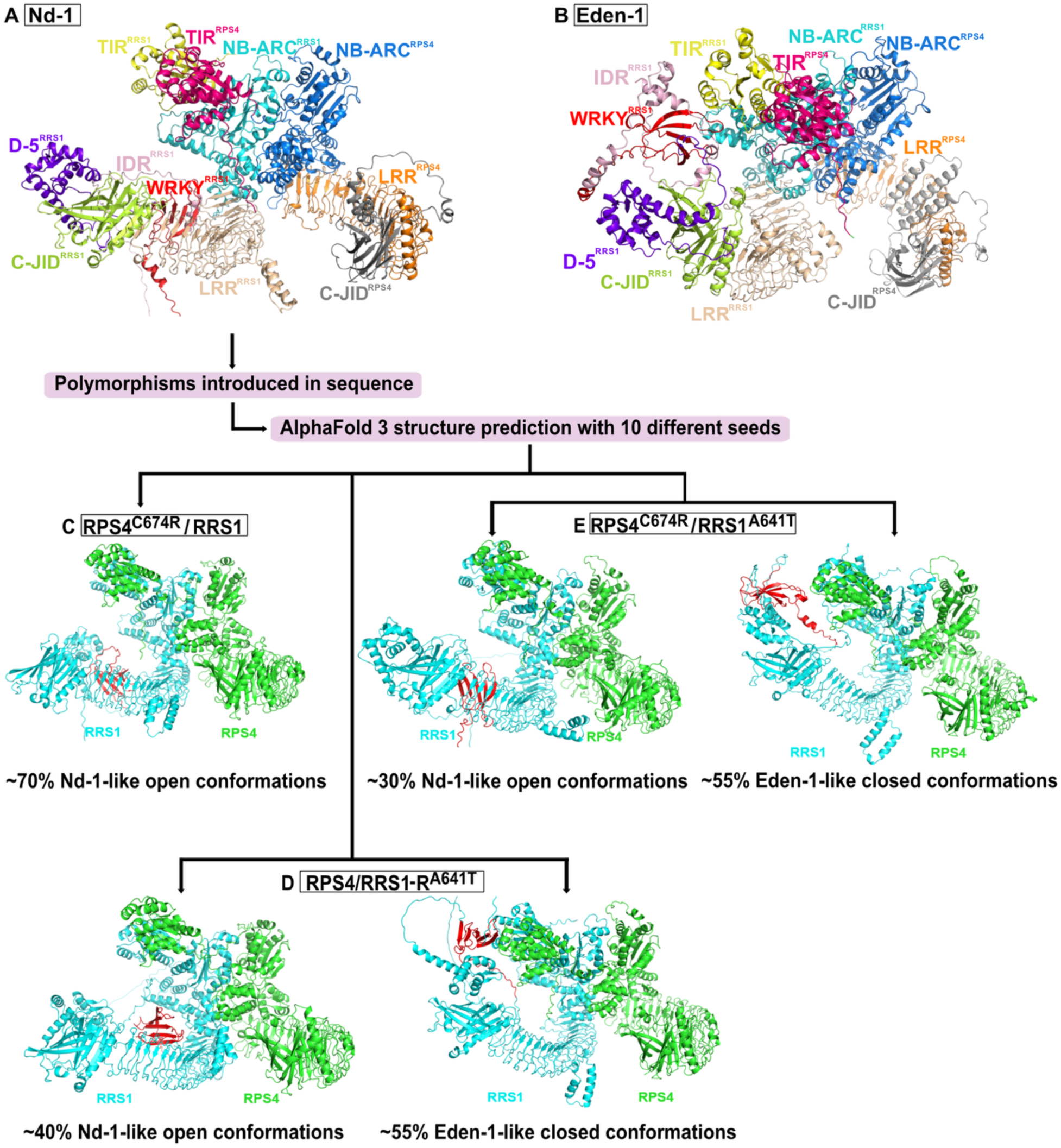
Mapping the non-synonymous polymorphisms in the Nd-1 sequence reveals insights on their significance in driving domain re-orientation. Most prominent representative conformation of AlphaFold 3-predicted structures of the RPS4/RRS1 heterodimeric complex from the accessions **(A)** Nd-1 and **(B)** Eden-1, in cartoon representation, with their domains labelled and coloured. **(C-E)** Structural predictions incorporating non-synonymous polymorphisms in the accession Nd-1: **(C)** RPS4^C674R^/RRS1 is observed to retain the Nd-1-like conformations. **(D)** RPS4/RRS1^A641T^ shows a noticeable increase in Eden-1-like conformations. **(E)** RPS4^C674R^/RRS1^A641T^ features both Nd-1-like and Eden-1-like conformations. Panels **(A)** and **(B)** display the Nd-1 and Eden-1 models respectively, with their RPS4/RRS1 chain domains highlighted in distinct colours. Each sub-panel represents the most frequent conformations among ten consecutive AlphaFold 3 predictions, obtained using different random seeds. Structural conformations of the different RPS4/RRS1 models are shown in cartoon representation with the RRS1’s WRKY domain coloured in red.

We then examined the potential roles of the key Eden-1-specific non-synonymous polymorphisms C674R (RPS4) and A641T (RRS1), by assessing their effect on the predicted conformational ensembles, when introduced within the RPS4 and RRS1 protein sequences of the Nd-1 accession (Fig. 3A). Noticeable shifts were observed in the distribution of conformations among Nd-1-like (LRR and adjacent C-terminal structured domains positioned away from TIR), Eden-1-like (LRR and adjacent C-terminal structured domains oriented towards RRS1 TIR), and alternative states. The C674R polymorphism largely preserved the Nd-1-like structural conformation, whereas A641T preferentially retained Eden-1-like conformations (Fig. 3C, D, Table S9). The C674R-A641T double substitution also predicted a substantial number of conformations similar to Eden-1, while also retaining a smaller proportion of models in the Nd-1-like conformation (Fig. 3E). Together, these findings predict a key role of the A641T polymorphism in RPS4/RRS1 complex structural dynamics and modulation of the RRS1’s structured C-terminal domains re-orientation, and consequently in the stability of the inactive complex.

To experimentally validate whether the naturally occurring A641T substitution in *RPS4/RRS1^Eden-1^* is responsible for thermostable disease resistance to GMI1000, we performed genetic complementation assays of the *rps4-21/rrs1-1* double knockout mutant with three different genomic constructs. Two independent T2 lines of each construct were selected for phenotypic characterization: i) the native *RPS4/RRS1^Nd-1^* haplotype (CNd-1#1, CNd-1#2), ii) the native *RPS4/RRS1^Eden-1^* haplotype (CEd-1#1, CEd-1#2), and iii) the *RPS4/RRS1^Nd-1^ ^A641T^* construct in which the A641T substitution from Eden-1 was introduced into the *RPS4/RRS1* genomic clone from Nd-1 (CA641T#1 and CA641T#2) (Fig 4A). At 27°C and 30°C, *RPS4* and *RRS1* expression levels measured at 4 dai were similar to, or higher than, those observed in Eden-1 (Fig. S5, Table S10). Along with the *rps4-21/rrs1-1* double mutant and Eden-1 accession, we monitored the phenotypic response of the complemented lines for disease response to GMI1000 at 27°C and 30°C. As expected, the *rps4-21/rrs1-1* mutant displayed wilting symptoms at both temperatures, while Eden-1 retained full resistance. The CNd-1#1 and CNd-1#2 lines were resistant at 27°C but became more susceptible at 30°C, in contrast to the CEd-1#1 and CEd-1#2 lines that maintained a thermostable disease resistance (Fig. 4A; Table S11). The CA641T#1 and CA641T#2 lines were resistant at 27°C, confirming the functionality of the *RPS4/RRS1^Nd-1^* ^A641T^ immune receptor complex in the complemented *rps4-21/rrs1-1* double mutant. However, CA641T#1 and CA641T#2 lines were only partially resistant at 30°C, compared to the CEd-1 complemented lines (Fig. 4A). The *rrs1-1* single mutant (*17*) was also complemented with either *RRS1^Nd-1^, RRS1^Eden-1^*, or *RRS1^Nd-1^ ^A641T^* genomic constructs. All transgenic lines were thermolabile suggesting that the complementation with *RRS1* alone is not sufficient to restore thermostability (Fig. S6A; Table S12). These data suggest that additional genetic elements present in the *RPS4/RRS1^Eden-1^* genomic clone, and likely in the *RPS4* gene from Eden-1, are required for immune receptor complex thermostability. Consistent with this observation, the introduction of the reverse substitution T641A in *RPS4/RRS1^Eden-1^* (hereafter designated as *RPS4/RRS1^Eden-1^ ^T641A^*) did not fully abrogate the thermostable immune response of the selected transgenic lines (Fig. S6B; Table S13).

**Fig. 4.**
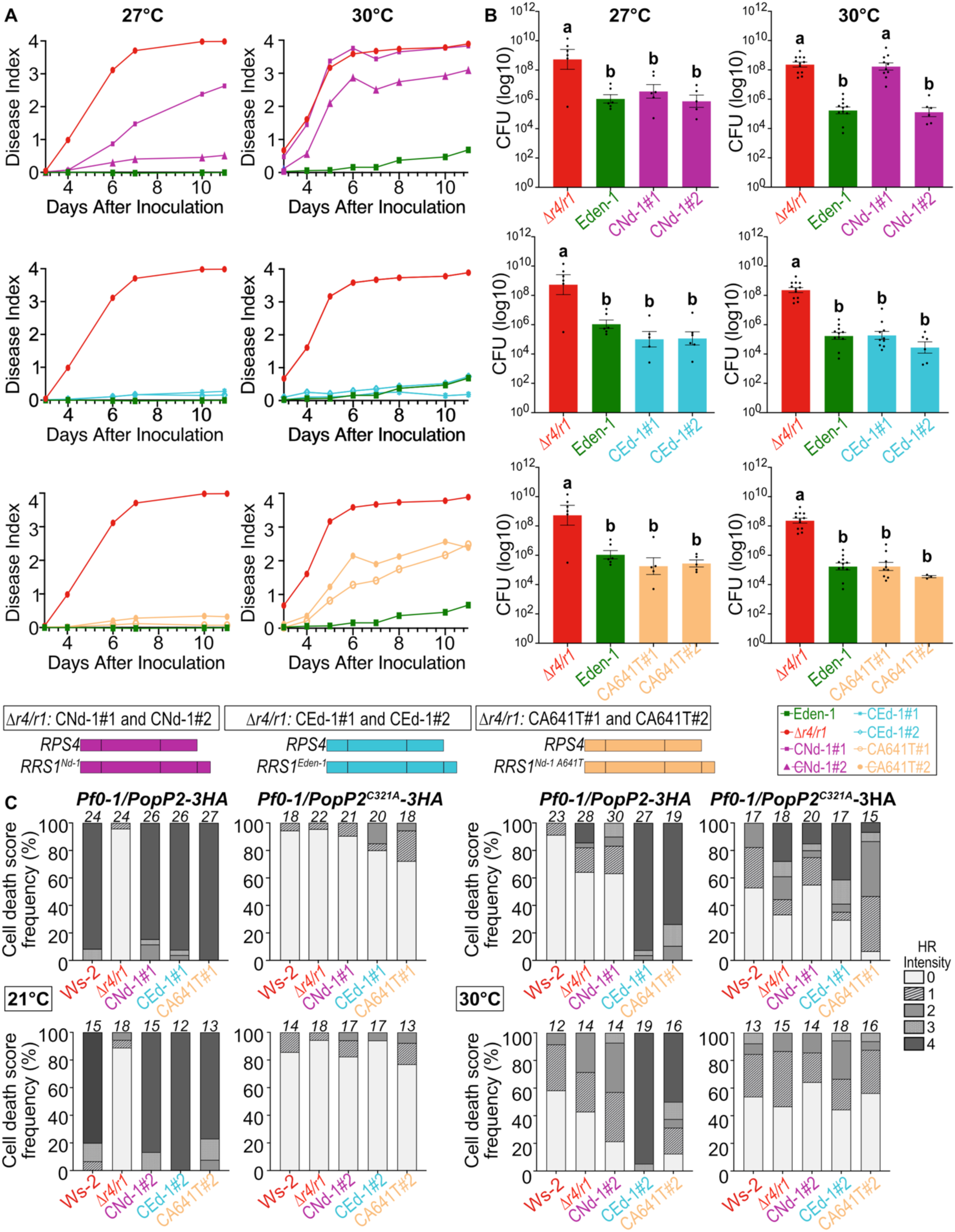
The *RPS4/RRS1^Eden-1^*haplotype confers thermostable resistance to GMI1000. The *rps4-21/rrs1-1* (Δ*r4/r1*) double mutant was transformed with three genomic constructs: CNd-1#1 and CNd-1#2 (native *RPS4/RRS1^Nd-1^*), CEd-1#1 and CEd-1#2 (native *RPS4/RRS1^Eden-1^*), and CA641T#1 and CA641T#2 (*RPS4/RRS1^Nd-1^ ^A641T^* with the A641T mutation). Two independent T2 lines per construct were selected for complementation assays. (**A**) Disease incidence following GMI1000 inoculation of complement lines and the *rps4-21/rrs1-1* double mutant at 27 °C and 30 °C. (**B**) Bacterial colonization assays at disease index of 1. Data represent log_10_ transformed colony-forming units (CFU) ± SEM (standard error of the mean). Tukey’s test groupings are indicated by lowercase letters. (**C**) Hypersensitive response assays at 21°C and 30°C using *Pseudomonas fluorescens* delivering active PopP2 or inactive PopP2^C321A^. Italicized numbers above bars indicate the number of infiltrated leaves. Cell-death symptoms are represented by rectangles whose color reflects the frequency of cell-death score frequency in infiltrated zones, ranging from absence (0, white) to complete cell death (4, black) of infiltrated zones.

We then assessed, at 4 dai, the expression of *RPS4, RRS1, SAG13* involved in disease response to bacterial infection (*12*), and the heat-responsive gene *HSP70* (*34*) at 27°C and 30°C. At 30°C, *HSP70* expression was strongly induced in all lines, while a decrease in expression was observed for the other genes (Fig. S5, Table S10). At 27°C, *RPS4*, *RRS1*, and *SAG13* were upregulated upon inoculation in all lines, except in the non-complemented susceptible *rps4-21/rrs1-1* double mutant. At 30°C, *RPS4*, *RRS1*, and *SAG13* remained upregulated upon inoculation in all lines, except for the susceptible *rps4-21/rrs1-1* mutant and the CNd-1#1 line, which exhibits an intermediate phenotype (Fig. S5). These results suggest that the *RPS4/RRS1^Eden-1^* complex not only contributes to pathogen recognition but also modulates plant responses to combined heat and biotic stress. Additionally, at 4 dai, disease symptoms of all tested transgenic lines were consistent with bacterial colonization levels (Fig. 4B; Table S14), supporting the activation of RPS4/RRS1-mediated immune response.

To further validate whether the thermostable induction of resistance mechanisms depends on the *RPS4/RRS1* pair, we conducted, at both 21°C and 30°C, a *Pseudomonas fluorescens* (*Pf0-1*)-mediated effector delivery assay, allowing us to monitor activation of the RPS4/RRS1 immune complex by wild-type PopP2, but not the PopP2^C321A^ catalytic mutant. Upon receptor pair activation, a strong cell-death response can be observed 24h after effector delivery in the leaves of 4-week-old plants carrying functional *RPS4/RRS1* alleles (*10*) (Figs. 4C, S7, S8, Table S15). At 21°C, except for the *rps4-21/rrs1-1* double mutant, Eden-1 and all complemented lines showed a strong cell-death response in the leaves following infiltration with *Pf0-1/PopP2-3HA* (Figs. 4C, S7; table S15), but not with *Pf0-1/PopP2^C321A^-3HA*, indicating proper recognition of PopP2 by the transgenic expression of the different NLR pair variants. At 30°C, transgenic lines carrying *RPS4/RRS1^Eden-1^* and *RPS4/RRS1^Nd-1^ ^A641T^*, but not *RPS4/RRS1^Nd-1^*, developed a cell-death response in the presence of PopP2. However, the response in CA641 complemented lines was more variable (Fig. 4C). These data indicate that both RPS4/RRS1^Eden-1^ and RPS4/RRS1^Nd-1^ ^A641T^ receptor complexes retain their sensing and signaling functions at elevated temperatures in adult plant leaves.

The exploration of genetic diversity within plant species, combined with genetic mapping, structural modelling, and functional genetics, could allow the identification of key polymorphisms associated with thermostability of resistance to pathogen infection. As previously observed for glyphosate resistance in weeds (*35*), and toxin resistance in butterflies (*36*), our data suggest that a synergistic combination of mutations in the natural allelic *RPS4/RRS1* variant of Eden-1 maintains resistance to pathogen infection under heat stress. In addition to the two Eden-1-specific amino-acid changes, we obtained further insights into the contributions from the other amino-acid changes in *RPS4/RRS1^Eden-1^* compared to Nd-1 (Fig. S9) than by introducing them into the RPS4/RRS1^Nd-1^ structure. They were predicted to have an overall destabilizing effect (Tables S16, S17) within the immediate local environment, as well as in terms of combined effects (Table S18), consistent with perturbing intramolecular contacts and cooperative networks that drive domain organization and lead to conformational changes. In terms of individual polymorphisms, the non-synonymous substitution A641T in the RRS1 LRR domain consistently showed the most noticeable effects (Tables S16, S17). However, we cannot rule out that these polymorphisms stabilize the activated resistosome structure directly. A major limitation in analyzing the structural basis of such effects is the paucity of experimental structural information for the RPS4/RRS1 pair. Currently, the only available structural information for NLRs derives from the resistosome structures of ROQ1 (*37*) and RPP1 (*38*). No structural data exist for the inactive state, though it is assumed to resemble the monomeric state described for the coiled-coil domain containing NLR ZAR1 (*39*). Given that NLRs are present in plants and mammals, structural modeling combined with the analysis of their natural allelic diversity among populations provides an opportunity to pinpoint polymorphisms that confer thermostable immune function and may help mitigate climate-driven disease outbreaks

Our study reinforces the importance of conserving and exploring natural genetic diversity, a reservoir of adaptive mechanisms against pathogens under abiotic stressors and more generally against multiple stresses. These resources offer a promising avenue to engineer resilient crops in an ever-changing environment, thereby contributing to global food security.

## Acknowledgments

We would like to thank Cédric Glorieux for generating the Eden-1 × *pi-1* cross. We thank Fernando Rab, Detlef Weigel, and his team for providing *Arabidopsis* seeds from the worldwide collection. We are grateful to Baptiste Mayjonade for his assistance and advice with the extraction of high-molecular-weight genomic DNA, and to Henri Desaint for contributing to the design of nonpolymorphic tailed primers used in the polymorphism analysis of the *RPS4/RRS1* locus. We also thank Ludovic Legrand for his assistance and for submitting the *RPS4/RRS1* locus sequence data and raw sequencing data from the eight BSA pools. We thank Adrien Belny for developing the R script used to compute LS means. We are also grateful to Qisheng Pan for his help with accessing and troubleshooting the DynaMut2 software. We humbly acknowledge the valuable inputs and software help provided by Dalton Ngu.

## Author contributions

Conceptualization: NA, BK, LD, FR, RB

Data curation: NA, MS, CJ, MN, BK, FR, RB

Analysis: NA, MS, CJ, MN, SC, JG, RD, MB, BK, LD, FR, RB

Funding acquisition: BK, LD, FR, RB

Experiments / Investigation: NA, MS, CG, ED, HD, MZ, MM, MB, LD, FR, RB

Methodology: NA, MS, LD, FR, RB

Project administration: NA, MB, BK, LD, FR, RB

Resources: NA, RD, AMH, MB, BK, LD, FR, RB

Software: NA, MS, SC, JG, RD, FR, RB

Supervision: MN, MB, BK, LD, FR, RB

Validation: NA, MS, MN, MB, BK, LD, FR, RB

Visualization: NA, MS, MN, MB, BK, LD, FR, RB

Writing – original draft: NA, LD, FR, RB

Writing – review & editing: NA, MS, CJ, MN, HD, SC, JG, RD, AMH, MB, BK, LD, FR, RB

## Data and materials availability

Except for HiFi sequence genomes from Eden-1 and *pi-1* genotypes newly generated in this study, all data are available in the main text or the supplementary materials. The Whole Genome Shotgun projects have been deposited at DDBJ/ENA/GenBank under the accession of JBPXLL000000000 and JBPZNV000000000 for Eden-1 and *pi-1* genotypes. PacBio sequences of *RPS4/RRS1* locus are publicly available under the https://doi.org/10.57745/LPMXI3. Raw sequencing data of the eight pools used in the BSA approach have been deposited at NCBI under the BioProject PRJNA1369404. RPS4/RRS1 sequences of Nd-1 and Eden-1 used for protein structure modeling are provided as fasta sequences.

## Supplementary Materials

Materials and Methods

Figs. S1 to S9

Tables S1 to S21

References (40–67)

## Materials and Methods

### Bacterial strains, plant materials, and growth and inoculation conditions

The *R. pseudosolanacearum* GMI1000 reference strain and mutated strain in *PopP2* (GRS100) were previously described (*12*) and used for inoculation experiments. These strains were grown at 28°C on either a complete BG solid or liquid medium (*40*) containing gentamicin (7.5 µg/mL) for GRS100. *Pseudomonas fluorescens* (*Pf0-1*) strains used, expressing either wild-type PopP2 or the catalytically inactive PopP2^C321A^ mutant (*12*), were grown on King’s B solid and liquid medium containing chloramphenicol (30 μg/mL), tetracycline (5 μg/mL), and gentamicin (15 μg/mL), at 28 °C.

The 103 accessions of *Arabidopsis thaliana* used in this study are listed in Table S1. The *rps4-21/rrs1-1* double mutant in the Ws-2 background (*17*) was used to generate all transgenic lines for complementation assays. All inoculation assays on natural accessions, the F2 population (N=1978) resulting from a cross between Eden-1 and an the *pistillata-1* (*pi-1*) EMS mutant in the Ler background (NASC stock number NW77, ABRC stock number CS77) (*41*), and the genetic complements were conducted following the root cut condition at 27°C and 30°C as described in Aoun et al., 2017 (*15*). Wilting symptoms were scored based on a scale from 0 (no wilting leaves) to 4 (all leaves are wilted) (*42*). Symptoms were monitored from 3 up to 14 days after inoculation (dai) (Tables S2-S4, S11-S13).

### Disease incidence and *A. thaliana* colonization assays

For each time point (dai) at each combination of temperature [27°C or 30°C], disease ratings were adjusted using the following linear model:

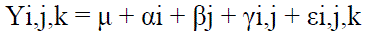

where Yi,j,k denotes the phenotype of the k^th^ plant of the j^th^ replicate of the i^th^ genotype; μ is the intercept; α is the genotypic fixed effect; β is the replicate fixed effect; γ is the replicate × genotype interaction fixed effects; and ε are the residuals with εi,j,k → N (0, σ2ε) independently and identically distributed. Least-square-means (LSmeans) of the genotypic effect and their associated confidence intervals (confidence level of 0.95) were obtained using the linear model (Tables S2, S11-S13). Contrast tests were made, allowing comparison between each pair of genotypes (Table S19). The R package *emmeans* (*43*) was used to compute LSmeans and to perform contrast tests. We performed colonization assays as described in Morel et al., 2018 (*44*). We used a Tukey HSD test to compare *in planta* bacterial multiplication relative to *the rps4-21/rrs1-1* double mutant.

### Genetic architecture of the resistance and bulk segregant analysis

To estimate the genetic architecture associated with the thermostable resistance of Eden-1, we adopted a bulk segregant analysis approach. We phenotyped 1,978 F2 progeny resulting from a cross between Eden-1 accession and the *pi-1* EMS mutant. One leaf of each labeled plant was sampled for DNA extraction and sequencing just before inoculation with the GMI1000 strain. The genomic DNA extraction protocol was adapted from QIAGEN DNeasy kit^®^ (Qiagen GmbH, Hilden, Germany). DNA samples were purified using guanidinium chloride (8M) and then treated with RNAse A (ribonuclease A) (Thermo Fisher Scientific). Purified DNA samples were quantified using the Quanti-iT^TM^ Picogreen® dsDNA kit (Thermo Fisher Scientific™) to compose equimolar DNA pools of 167 individuals each and combined to form one pool of resistant and seven pools of susceptible F2 plants. DNA sequencing was performed on Illumina HiSeq3000 using libraries prepared with TruSeq Nano DNA sample kits (Illumina). Pooled libraries were subjected to 150-pb paired-end sequencing. Demultiplexed FastQ files were generated using the bcl2fastq conversion software v2.17.

*De novo* genomes for the parental lines Eden-1 (JBPXLL000000000) and *pi-1* (JBPZNV000000000) were obtained by using the PacBio Single Molecule Real-Time technology. The SMRTBell library was prepared using a SMRTbell Express Template prep kit, following the “procedure and checklist-preparing > 30 kb libraries using SMRTbell Express Template preparation kit” protocol. Genomic DNA (5 μg) was sheared with the 75 kb program using a Diagenode Megaruptor (Diagenode), generating DNA fragments of approximately 25kb. A Femto Pulse (Agilent Technologies, Santa Clara, CA, USA) assay was used to assess the fragment size distribution. Sheared genomic DNA was carried into the first enzymatic reaction to repair any damage that may be present on the DNA backbone. A-tailing reaction followed by the overhang adapter ligation was conducted to generate the SMRT Bell template. After a 0.6 X AMPure PB beads purification, the samples were size selected using the BluePippin (Sage Science, Beverly, MA, USA) to recover all the material above 20 kb. The eluted DNA was then purified with 1X AMPure PB Beads to obtain the final libraries around 25 kb. The SMRTBell libraries were quality inspected and quantified on a Femto Pulse (Agilent Technologies) and a Qubit fluorimeter with Qubit dsDNA HS reagent Assay kit (Life Technologies). A ready-to-sequence SMRTBell Polymerase Complex was created using a binding Kit 2.1 (PacBio) and the primer V4. The diffusion loading protocol was used, according to the manufacturer’s instructions. The PacBio Sequel instrument was programmed to load and sequence the sample on PacBio SMRT cells v2.0 (Pacific Biosciences, Menlo Park, CA, USA), acquiring one movie of 600 min per SMRTcell. The work was performed at the Gentyane Sequencing Platform (Clermont-Ferrand, France).

For genome assembly of each parental line, reads were corrected with CANU 1.8 and used as inputs in SMARTdenovo (Jue Ruan, https://github.com/ruanjue/smartdenovo) and wtdbg2.0 (*45*) software. The first wtdbg2.0 assembly was performed by using corrected reads only, while the second wtdbg2.0 assembly was performed by using both corrected reads and raw reads. The sequences of these four primary assemblies were transformed into pseudo-long reads of 100 kb with an overlap of 50 kb. Then, the pseudo-long reads longer than 20 kb were assembled with CANU 1.8 (–trim-assemble –PacBio-corrected). After removal of spurious contigs (i.e., short contigs included in longer contigs; overlaps detected by minimap2-x asm5 (*46*), the pseudo-chromosomes were obtained with ALLMAPS (*47*) by scaffolding the contigs using the *A. thaliana* Col-0 genome (TAIR Arabidopsis) as a reference. Overlaps detected by minimap2-x asm5 were transformed into pseudo-genetic markers (∼1 marker per 30kb). Two iterations of polishing with arrows were performed on the raw genome sequences, including mitochondrial and chloroplast genomes. Protein-coding genes were annotated by integrating five sources of information. Priority was successively given to: i) a BLASTp search of reciprocal best hits with 15,832 “taxonomy:3702” (*A. thaliana*) proteins tagged as reviewed in the Uniprot database (90% span, 80% identity) as of March 2019 (Uniprot C), ii) EC numbers assigned by using the protocol (E2P2 v3.1, (*48*) with BLAST e-value cutoff lowered to 1.e-5 and pathway-prediction-score set to 0.3 in pathway-tools, iii) the transcription factors and kinases identified by ITAK release 1.7 (*49*), iv) the transcription factors identified by PlantTFCat (*50*), and v) the Interpro (release 72.0) search (*51*). The EC numbers were tested against the ENZYME (*52*) database (release March 2019), updated when deprecated and then used to get the description of the enzymes. Finally, the protein annotations were validated by the tbl2asn software (https://www.ncbi.nlm.nih.gov/genbank/tbl2asn, March 2019). GO terms were assigned using the BLAST2GO pro software, integrating blastp similarities with Genbank NR (*53*) (release March 2019, e-value < 1e-5, 20 best-scoring hits) and Interpro (release 72.0) results.

We performed sequence analysis of F2 plants on the Galaxy platform using the bwa mapping software (*54*) and following the “SNP detection and effect prediction” protocol by using the two *de novo* assembly genomes from the Eden-1 accession and the *pi-1* mutant. The depth of sequencing of the eight pools was, on average, 51X (min: 43X, max: 55X) when mapped on either Eden-1 or *pi-1*. We mapped raw reads of each of the eight pools onto the two reference genomes with a maximum of four mismatches on at least 80 nucleotides. For SNP calling, the minimum position coverage of supporting reads was 20, the minimum variant coverage was 10, the minimum variant frequency was 0.2, and the minimum homozygous frequency was 0.75. Following the analysis, the allele read count matrix, for the reference and alternate alleles, was composed of 354,763 and 316,504 SNPs across the eight pools using Eden-1 and *pi-1* reference genomes, respectively. We filtered the output for i) non-nuclear SNPs, ii) for SNPs without mapped reads in at least two pools out of the seven pools of susceptible F2 plants, and iii) for missing data corresponding to unmapped genomic positions (number of remaining SNPs with Eden-1 and *pi-1* reference genomes after filtering: 341,094 and 313,178 SNPs, respectively). Then we filtered our data according to a distribution of the depth of raw reads per SNP obtained for each pool, and we removed SNPs with extreme values. For resistance and susceptible pools, we kept SNPs for which the depth of reads ranged between 26 and 65 (number of remaining SNPs with Eden-1 and *pi-1* reference genomes: 285,873 and 253,884 SNPs, respectively). Finally, to identify QTL regions underlying thermostable resistance of Eden-1, we calculated for each SNP the difference between the reference allele frequency obtained in the resistant pool and the average of the reference allele frequencies between the seven susceptible pools, hereafter named ΔSNP. Then, to aggregate ΔSNP results among physically linked SNPs, we calculated at each focal SNP position an average ΔSNP based on all the SNPs present in a 500kb window centered on that focal SNP. To illustrate our results, plots were computed using the “ggplot2” package (*55*) under the *R* environment 3.6.1 (*56*).

### Loci cloning and genetic complementation

To generate the genetic constructs used for transgenic complementation we used Gateway^TM^ technology (Invitrogen, Carlsbad, CA, USA). The *RPS4/RRS1* genomic region from Nd-1 was chosen because it does code for thermolabile RPS4/RRS1 variants at 30°C and has the lowest numbers of polymorphisms when compared to that of Eden-1. Genomic DNA was extracted from Nd-1 and Eden-1 using a CTAB-based (hexadecyl trimethyl ammonium bromide; C19H42BrN) DNA extraction protocol adapted from Allen et al. (2006) (*57*). The corresponding *RPS4/RRS1* genomic region from both accessions, including the intergenic region as well as *RPS4* and *RRS1* terminator sequences, was PCR-amplified with ATT-GATE1 and ATT-GATE2 primers (Table S20) using PrimeStar HS DNA polymerase from Takara Bio Inc. (Otsu, Japan). Amplicons were purified with a QIAEX II Gel Extraction Kit (Qiagen GmbH, Hilden, Germany) and cloned into the pDONR207 vector using BP Clonase according to the manufacturer’s instructions (Invitrogen). The corresponding pENTR clones (Table S20) were verified by SANGER sequencing. Generated sequences were aligned on Clone Manager©.

We aligned the nucleotide sequences of *RPS4/RRS1^Eden-1^* with the long-read *RPS4/RRS1* sequences of 114 *A. thaliana* accessions (Table S1) using Bioedit 7.7.1.0 (*58*), and looked for sequence polymorphisms (Fig. 2A). Several polymorphisms specific of Eden-1 were identified, including amino-acid changes C674R and A641T in the LRR domains of RPS4 and RRS1, respectively. Using primers listed in Table S20 and PrimeStar HS DNA polymerase from Takara Bio Inc. (Otsu, Japan), a two-step PCR-based site-directed mutagenesis on the *RRS1^Nd-1^* genomic sequence was performed to generate the *RRS1^A641T^* variant. The pB7-RPS4-RRS1^Nd-1^ ^A641T^ binary vector was obtained by ligating a *Kpn*I-*Bam*HI restriction fragment containing the A641T mutation into a *Kpn*I-*Bam*HI-digested pENTR-RPS4/RRS1^Nd-1^ genomic clone (*12*).

As a result, we generated two binary vectors. The *RPS4/RRS1^Eden-1^*genomic fragment of the pENTR-RPS4/RRS1^Eden-1^ was transferred into the pB7-D35S-GWY-GFP binary vector using the LR Gateway reaction according to the manufacturer’s instructions (Invitrogen), resulting in pB7-RPS4-RRS1^Eden-1^. The pENTR-RPS4-RRS1^A641T^ plasmid was used to generate the pB7-RPS4-RRS1^A641T^. The binary vector pB7-RPS4-RRS1^Nd-1^ containing *RPS4/RRS1^Nd-1^* locus was already available (*42*). For complementation, all constructs were introduced in the *Agrobacterium tumefaciens* GV3101 strain and used for floral dip transformation (*59*) of five-week-old *A. thaliana rps4-21/rrs1-1* mutant. T1 and T2 progenies were selected using phosphinothricin (Duchefa) (7.5 μg/mL) and genotyped by PCR (Table S20).

Following this cloning strategy, we generated complemented transgenic lines in *rps4-21*/*rrs1-1* carrying the *RPS4/RRS1* sequences of Nd-1 (*RPS4/RRS1^Nd-1^*) (Nd-1#1-2), and Eden-1 (*RPS4/RRS1-1^Eden-1^*) (CEd-1#1-2). We also included a *RPS4/RRS1^Nd-1^* genomic clone in which the A641T polymorphism of Eden-1 RRS1 was introduced (*RPS4/RRS1^Nd-1^ ^A641T^*) (CA641T#1-2), and a *RPS4/RRS1^Eden-1^* genomic clone in which the T641A polymorphism of Nd-1 RRS1 was introduced (CT641A#1-2). We also complemented the *rrs1-1* single knockout mutant (*13*) with a genomic fragment restricted to *RRS1^Nd-1^* (CR1Nd-1#1-2), *RRS1^Eden-1^* (CR1Ed-1#3-6), and *RRS1^A641T^* (CR1A641T#1-4) alleles. We selected at least two independent transgenic lines among basta-resistant T2 progenies upon *Pfo-1*-mediated delivery of PopP2 in leaves at 23°C and hypersensitive response induction to ensure their functional complementation. Plasmid constructs, vector backbones, and purpose of use are presented in Table S21.

These transgenic lines were cultivated for seed production in greenhouse conditions (26.5°C +/-1.5°C, 16 h light). For selecting *A. thaliana* transgenic lines, seeds were sterilized as described in Morel et al. 2018 (*44*). Seeds were sown on MS medium supplemented with phosphinothricin (Duchefa) (7,5 μg/mL) for complemented lines of progeny selection. One week later, seedlings were transferred to Jiffy pots (Jiffy Products International AS, Norway) and grown for three more weeks for inoculation assays or hypersensitive response as described in Aoun et al., 2017 (*15*), and Tasset et al., 2010 (*10*), respectively.

### Hypersensitive response assays

For cell-death assays, we used *Pseudomonas fluorescens Pfo-1* strains delivering 3HA-tagged active PopP2 or inactive PopP2^C321A^ (*12*). We grew four-week-old plants as described above and then moved them inside climatic chambers (Memmert ®) set at 21°C or 30°C (8 h photoperiod) for 48 h prior *Pfo-1* infiltration for acclimation. *Pfo-1* strains were streaked out on King’s B agar plates containing appropriate antibiotics (Tetracycline 5 µg/ml, chloramphenicol 30 µg/ml for empty *Pfo-1* strain, additional gentamicin 15 µg/ml was added for *Pfo-1* carrying *PopP2-3HA* or *PopP2^C321A^-3HA* constructs) and grown overnight at 28°C. Pelleted bacteria were resuspended in 10 mM MgCl2 and the bacterial density was adjusted to OD_600nm_=0.2. Plant leaves were infiltrated with bacterial suspension using a 1 ml needleless syringe. At 6 h post-infiltration, samples were collected from three different plants (eight 4 mm^2^ discs) for immunoblot analyses. Cell-death symptoms were scored at 24 h after infiltration following a scale from 0 (no symptom) to 4 (complete necrosis).

### Immunoblot analyses

For PopP2-3HA or PopP2C321A-3HA detection, total protein extraction from eight 4 mm^2^ discs collected on three different plants at 6h post-infiltration for each temperature treatment was performed. Plant tissues were ground and resuspended in 100µl Laemmli buffer (0.125 M Tris HCl pH7.5, 4% SDS, 20% glycerol, 0.2M DTT) for western-blot analyses. After denaturation at 95°C for 3 min, samples were then loaded on sodium dodecyl sulfate polyacrylamide gels for electrophoresis and transferred to nitrocellulose membranes, which were blocked with 5% skimmed milk. Membranes were stained using Ponceau red staining to verify total protein loading. Membranes were then probed with the primary anti-HA-HRP antibody (3F10; Roche; dilution 1:5000). Protein detection was performed using the Bio-Rad Chemidoc Imaging system with the Bio-Rad Clarity Max Western ECL Substrate (Fig. S8).

### Gene expression analysis by RT-qPCR

We inoculated four-week-old plants with the GMI1000 strain and scored them for disease wilting as described above. At each temperature treatment and for inoculated and non-inoculated plants, three technical samples composed of two pooled leaves from two different plants were harvested from two biological replicates. This sampling took place when the disease score of *rps4-21/rrs1-1* double mutant plants, used as control, was 1. Total RNA was extracted using the Macherey-Nagel Nucleospin RNA Plus kit (New England Biolabs®), following the manufacturer’s instructions. We treated 5 µg of total RNA with DNase using the Invitrogen™ Ambion™ TURBO DNA-free Kit (Thermo Fisher Scientific©). cDNA was then generated with 1 µg of DNase-treated total RNA and Moloney Murine Leukemia Virus Reverse Transcriptase (M-MLV RT) (Life Technologies) according to manufacturer instructions. For qPCR reactions, we prepared mix with SYBR green (Takyon™ No ROX SYBR 2X Master Mix blue dTTP, Eurogentec©) and ran it on the Bio-Rad CFX Opus 384 Real-Time PCR instrument. For all genes of interest, 2 µL of cDNA reaction diluted 1:10 was used with gene-specific primers (Table S20) and presented as relative expression (2^-ΔCt^) to the mean Ct value of *At5G15710* (galactose oxidase/kelch repeat superfamily protein) and *At1G13320* (protein phosphatase 2A subunit A3) genes used as internal controls.

### Polymorphism analysis of *RPS4/RRS1* locus

Using Jal View 2.11.4.1 (*60*), we analyzed DNA polymorphism at the *RPS4/RRS1* locus of the two clones of *RPS4/RRS1^Eden-1^*and high-quality sequences of 114 *A. thaliana* natural accessions that we screened for their disease response to GMI1000 (Table S2). Sequences were obtained using two strategies: i) Consensus sequences were retrieved for each amplicon. Sequences of *RPS4* and *RRS*1 genes as well as corresponding to *RPS4/RRS1^Eden-1^*, were extracted using a custom Python script (v 3.9.0) from genome assemblies of 70 *A. thaliana* accessions (*61*, *62*) belonging to the panel used in this study, For each gene, homologous sequences were searched using the blastn program of Blast+ (v 2.15.0+) and aligned using MAFFT (v 7.453). ii) PacBio sequencing of two amplicons overlapping the *RPS4/RRS1* locus of 49 accessions phenotyped in our previous study (*15*) (Table S1). Amplicons were PCR amplified using nonpolymorphic tailed primers TRPS4end_F / TRRS1start_R and TRRS1start_F/ TRRS1end_R, designed by aligning Illumina sequences of *RPS4/RRS1* locus available in the 1001 genome project (*63*) for 124 out of 176 panel of accessions phenotyped in our previous study (*15*) (Table S20). PCR amplification and purification were performed as described above. SMRTbell® libraries using PacBio® barcoded universal primers for multiplexing were generated with PCR products mixed in equimolar quantities. Product purifications and pool for sequencing in MRTCell with a PacBio Sequel II sequencing system were realized at the Gentyane Plateforme (INRAE, Clermont-Ferrand, France). Consensus sequences were retrieved for each amplicon.

### Protein Structure Modelling

AlphaFold 3 (*24*) was chosen to perform structure predictions owing to its superior prediction accuracy, faster performance and its ability to predict larger, more complex biomolecular assemblies, leveraging an improved unified diffusion-based architecture that outperforms AlphaFold 2 in structural precision, conformational sampling, and multimer prediction (*64*). However, this tool does not allow us to predict structures comprising over 5000 tokens (1 token = 1 standard amino acid residue of protein, one atom of ligand, or one input nucleotide base), due to restraints on the size of the protein sequence allowed as input, the correlation of size with increasing computational requirements and decreasing quality of predictions, and the lack of training data for large structures. Therefore, we were able to predict only the RPS4/RRS1 heterodimer structures.

Ten sets of predicted models were generated for each sequence, resulting in 50 models for each accession. The ‘seed number’ for each prediction job was randomised, and template settings were ‘truned off’ to remove potential bias and let AlphaFold 3 capture diverse conformational space. For a conformation to be considered significant, we required at least 40% of the predictions to have Cα RMSD for structured domain regions <4 Å.

The predicted models were then evaluated using the following information: (i) pLDDT (predicted local distance difference test) and pTM (predicted template modelling) scores, and PAE (predicted alignment error) plots reported by AlphaFold 3; and (ii) structure superpositions of different predicted and experimental structures, analysing different interfaces, steric clashes and implausible conformations.

The overall pLDDT confidence values for most predicted structures were within the range expected for accurately predicted models, with higher confidence values for the core regions (70–90), and lower values (40–70) for flexible regions, with the lowest values (<40) predicted for C-terminal IDR stretches. The predicted structure of RPS4/RRS1^Nd-1^, coloured by pLDDT values, is shown in Fig. S3.

The pTM scores for screened models were also found to be consistently >0.5, indicating reasonable confidence in the prediction of the accurate fold, while their PAE plots indicated reasonable structural correlations between subunits and consistent heterodimer assembly (Fig. S4).

MutationExplorer (*33*) was used to infer the overall energy profiles of the predicted structures, which utilises Rosetta energy units (REU) as a relative scale used within the Rosetta framework to estimate the relative energies of protein conformations. Before MutationExplorer analysis, the selected structures were first energy-minimized (all-atom energy minimization through iterative cycles of side-chain repacking and backbone relaxation), using the Relax module of the ROSIE (Rosetta Online Server that Includes Everyone) server (*65*, *66*). This step helped resolve minor steric clashes, relax strained geometries and optimise hydrogen-bonding networks that can arise inadvertently from computationally generated structural models, thus ensuring the models occupy a physically realistic local energy minimum and are suitable for downstream structural analyses. Absolute energies estimated by using MutationExplorer corresponded to –8114.09 and –7909.91 REU (Rosetta energy units) for RPS4/RRS1^Eden-1^ and RPS4/RRS1^Nd-1^, respectively, suggesting higher stability for RPS4/RRS1^Eden-1^.

The DynaMut2 web server-based tool (*67*) was used to assess the stability contribution, as the difference in change in Gibbs free energy (ΔΔG), for different polymorphisms. This tool predicts the impact of point mutations on protein structure stability by integrating graph-based signatures with normal mode analysis (NMA), incorporating a knowledge-based potential energy function to estimate changes in ΔG. The introduction of Eden-1-specific polymorphisms into the RPS4/RRS1^Nd-1^ structure was predicted to have an overall destabilizing effect, especially pronounced for the polymorphisms within the RRS1 in its conformation at the local level (Tables S17, S18), consistent with the predicted domain reorganization when it changes conformation.

**Fig. S1.**
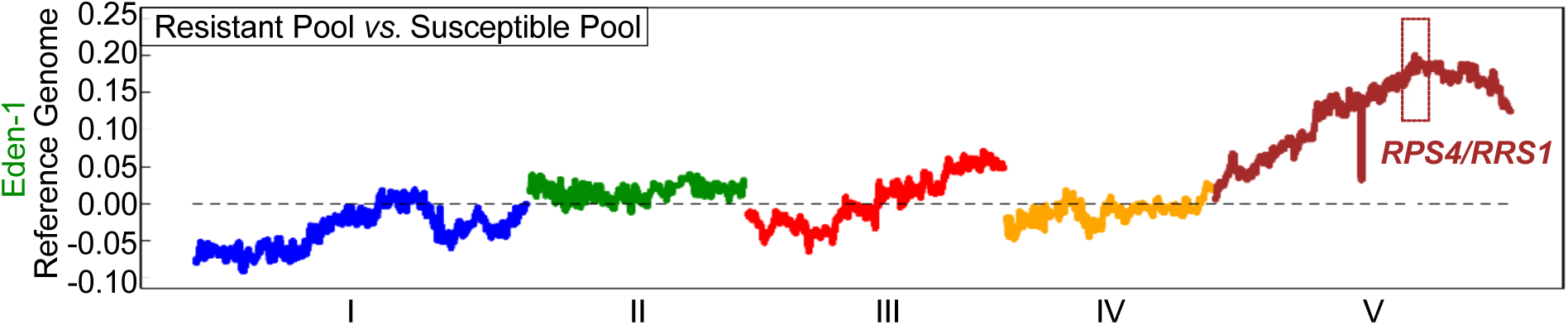
The *RPS4/RRS1* locus is genetically associated with *A. thaliana* disease response to GMI1000 at 30°C. Allele frequency differentiation analysis between resistant and susceptible pools identified a QTL on the Eden-1 genome, with the top associated SNP located 62 kb downstream of the RPS4/RRS1 locus. The x-axis corresponds to *A. thaliana* chromosomes, and the y-axis corresponds to allele frequency differences.

**Fig. S2.**
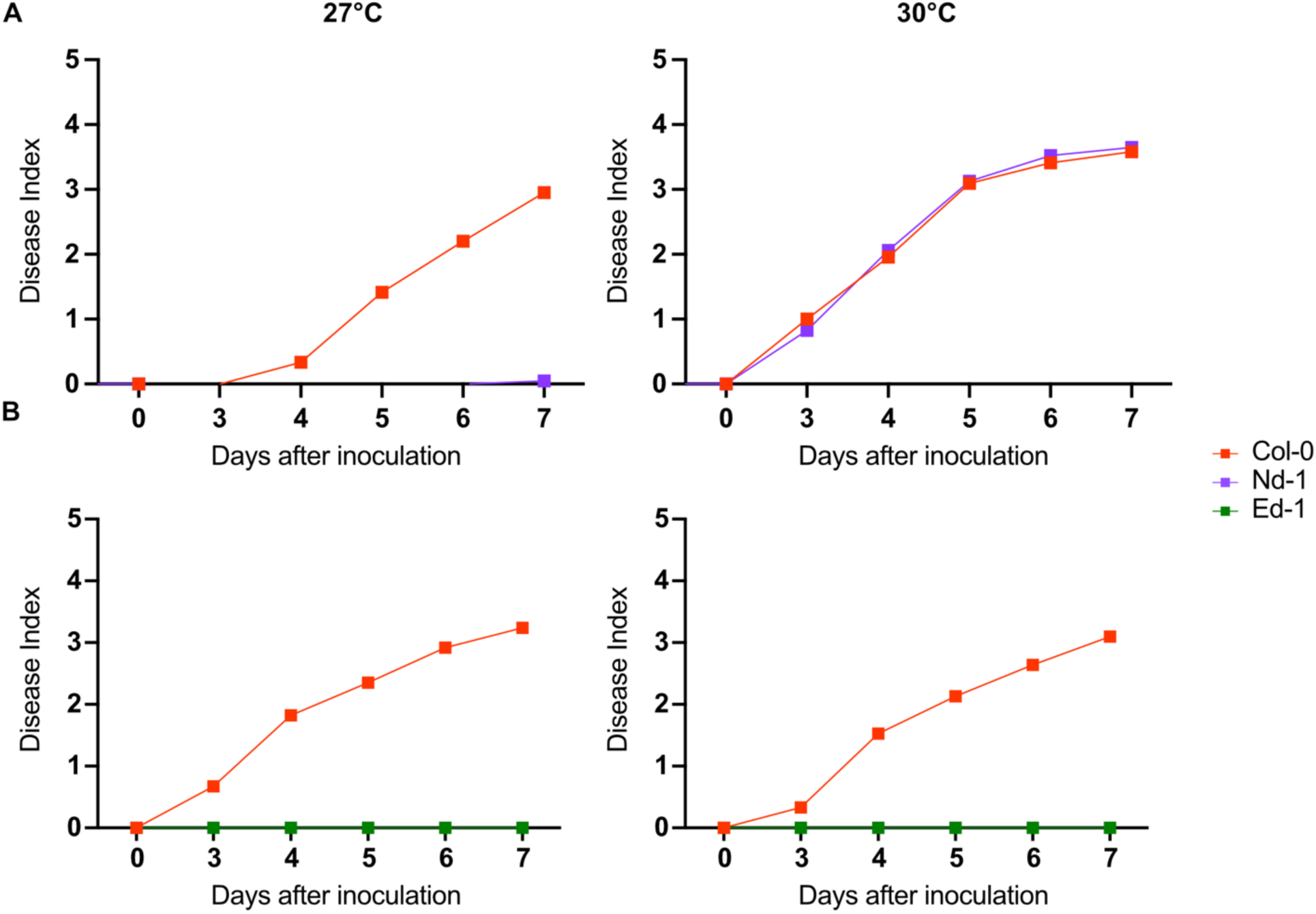
Disease wilting kinetic of Eden-1, Nd-1, and Col-0 plants against GMI1000 at 27°C and 30°C. (**A**) The thermolabile Nd-1 accession was tested along with susceptible Col-0 plants for disease response to GMI1000 at 27°C (left) and 30°C (right). Col-0 (N=43 plants) and Nd-1 (N=41 to 43 plants) were screened in two biological replicates at both temperatures with the GMI1000 strain. (**B**) Col-0 (N=72 plants) and Eden-1 (N=18 plants) were tested against GMI1000 at 27°C and 30°C. This experiment was repeated in three biological replicates. LS means were computed under the *R* environment.

**Fig. S3.**
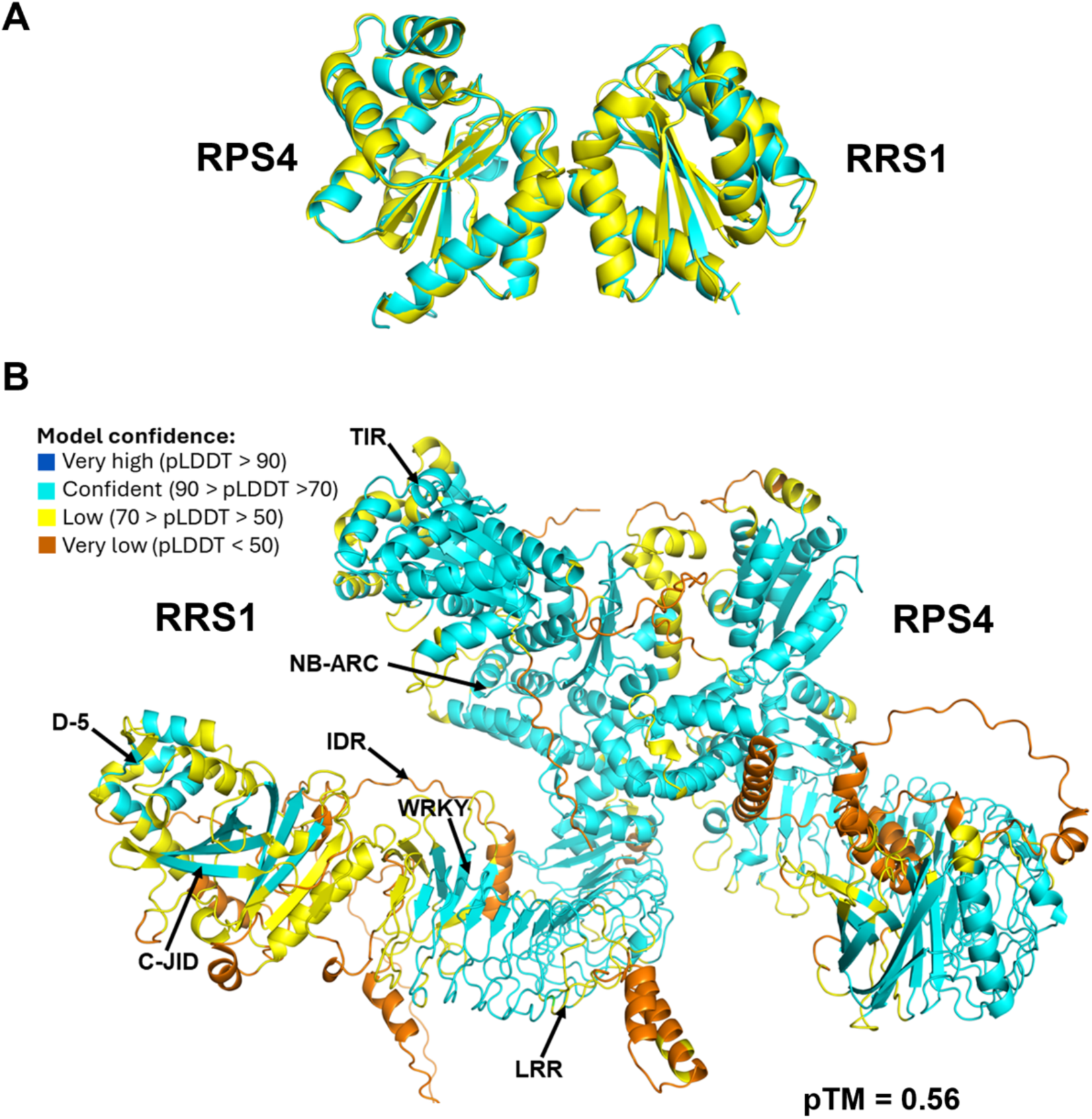
AlphaFold 3 structural prediction of the RPS4/RRS1^Nd-1^ TIR complex compared with the *A. thaliana* RPS4/RRS1 TIR domain complex crystal structure. (A) Structural superimposition of *A. thaliana* RPS4 TIR domain/RRS1 TIR domain complex (PDB: 4C6T) (cartoon, yellow) and AlphaFold 3-generated RPS4/RRS1^Nd-1^ TIR domain region (cartoon, cyan)**. (B)** AlphaFold 3-generated top-model for RPS4/RRS1^Nd-1^, colored by pLDDT values and highlighting the domains in RRS1.

**Fig. S4.**
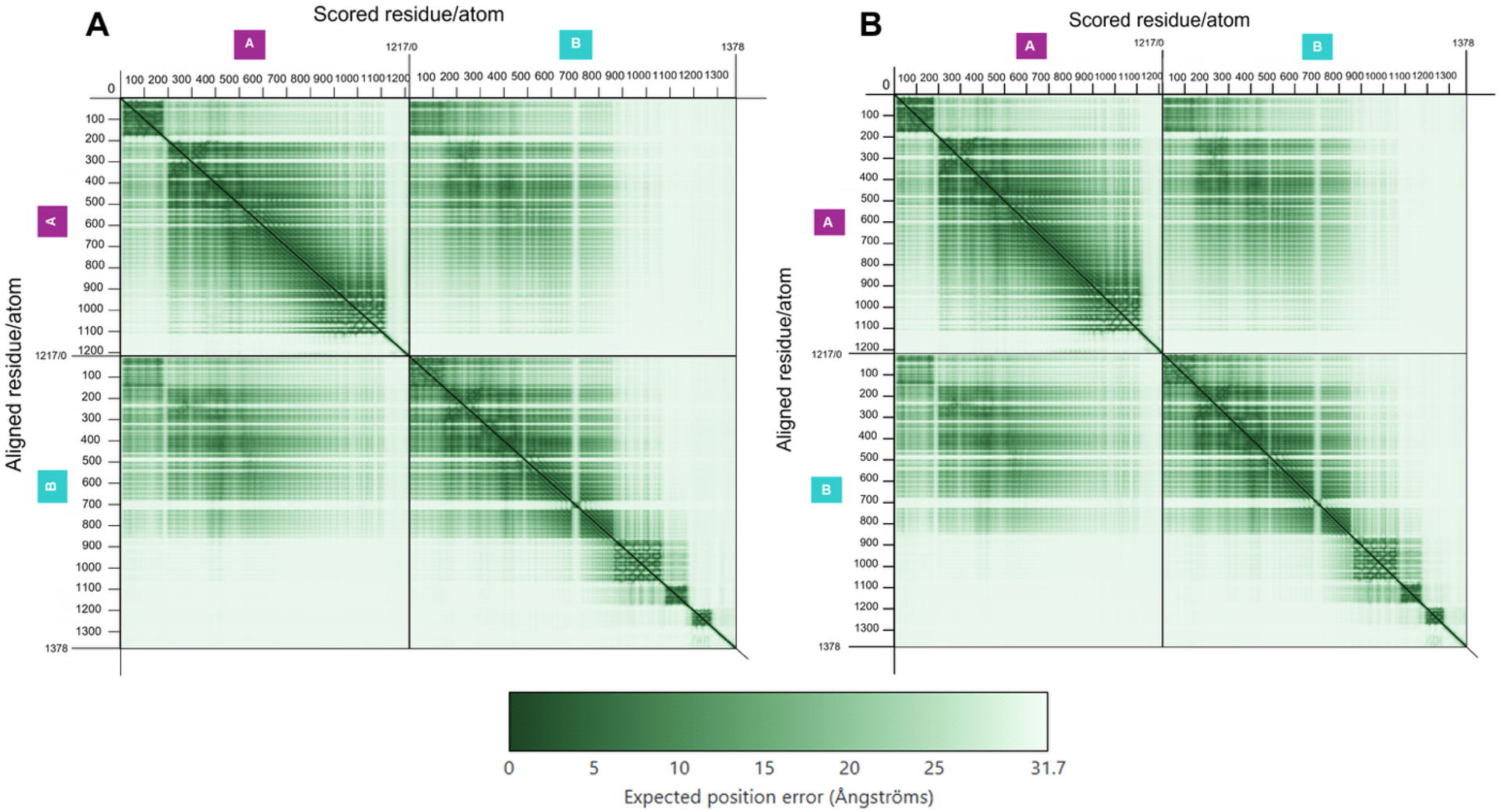
PAE plots for the selected conformations of AlphaFold 3-predicted RPS4/RRS1 structures for accessions Nd-1 (A) and Eden-1 (B). PAE plots indicate reasonable structural interactions between subunits and consistent heterodimer assembly in all the predicted models. The plots were generated using the PAE viewer web server (https://doi.org/10.1093/nar/gkad350), based on AlphaFold 3-derived .cif structure files and corresponding .json outputs for each model.

**Fig. S5.**
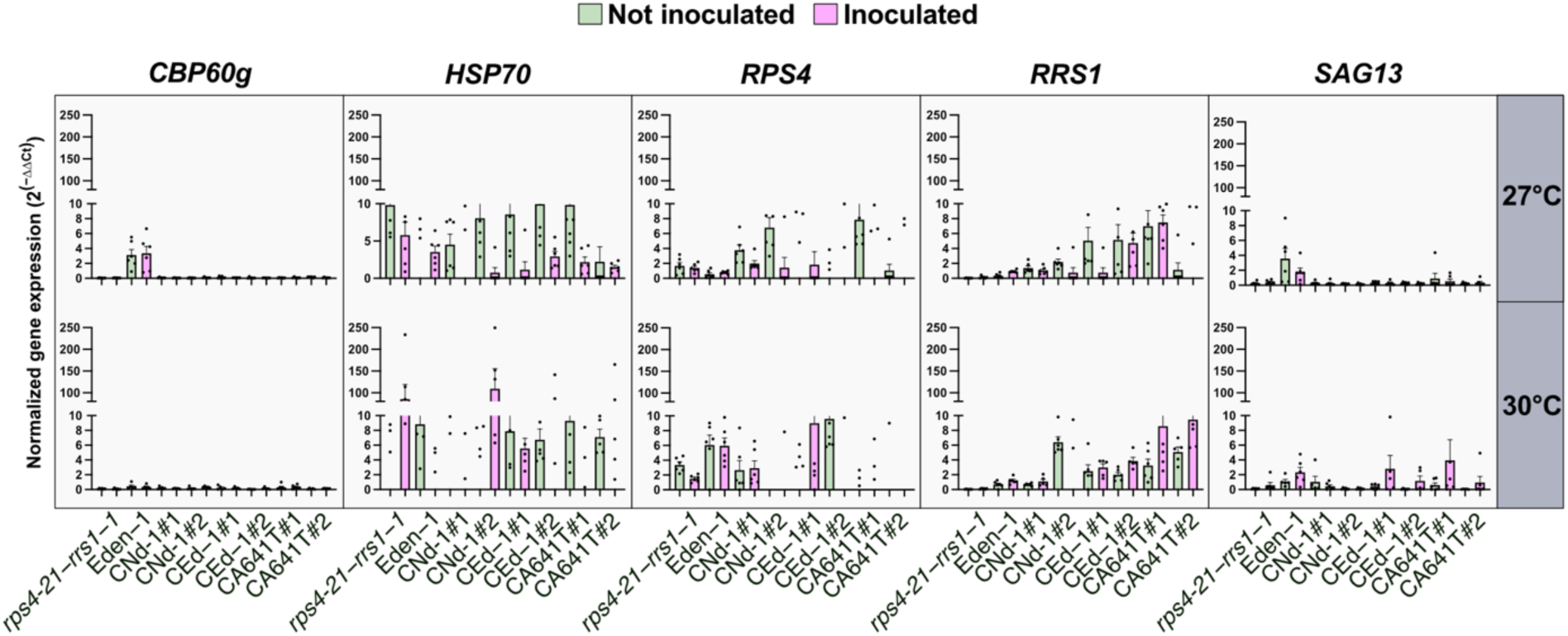
Relative gene expression analysis by RT-qPCR in genetically complemented plants inoculated with GMI1000 at 27°C and 30°C. Expression levels of *HSP70*, *RPS4*, *RRS1*, and *SAG13* were quantified in inoculated and non-inoculated plants at the time point when the disease score of the *rps4-21/rrs1-1* double mutant control reached 1. *CBP60g* has been previously described not be expressed at 28°C (*59*) and has been used as a control in this study. Relative expression was calculated using the 2^-ΔCt^ method, normalized to the mean Ct values of *At5G15710* and *At1G13320*, which served as internal reference genes. The data represent two biological replicates with three technical replicates each.

**Fig. S6.**
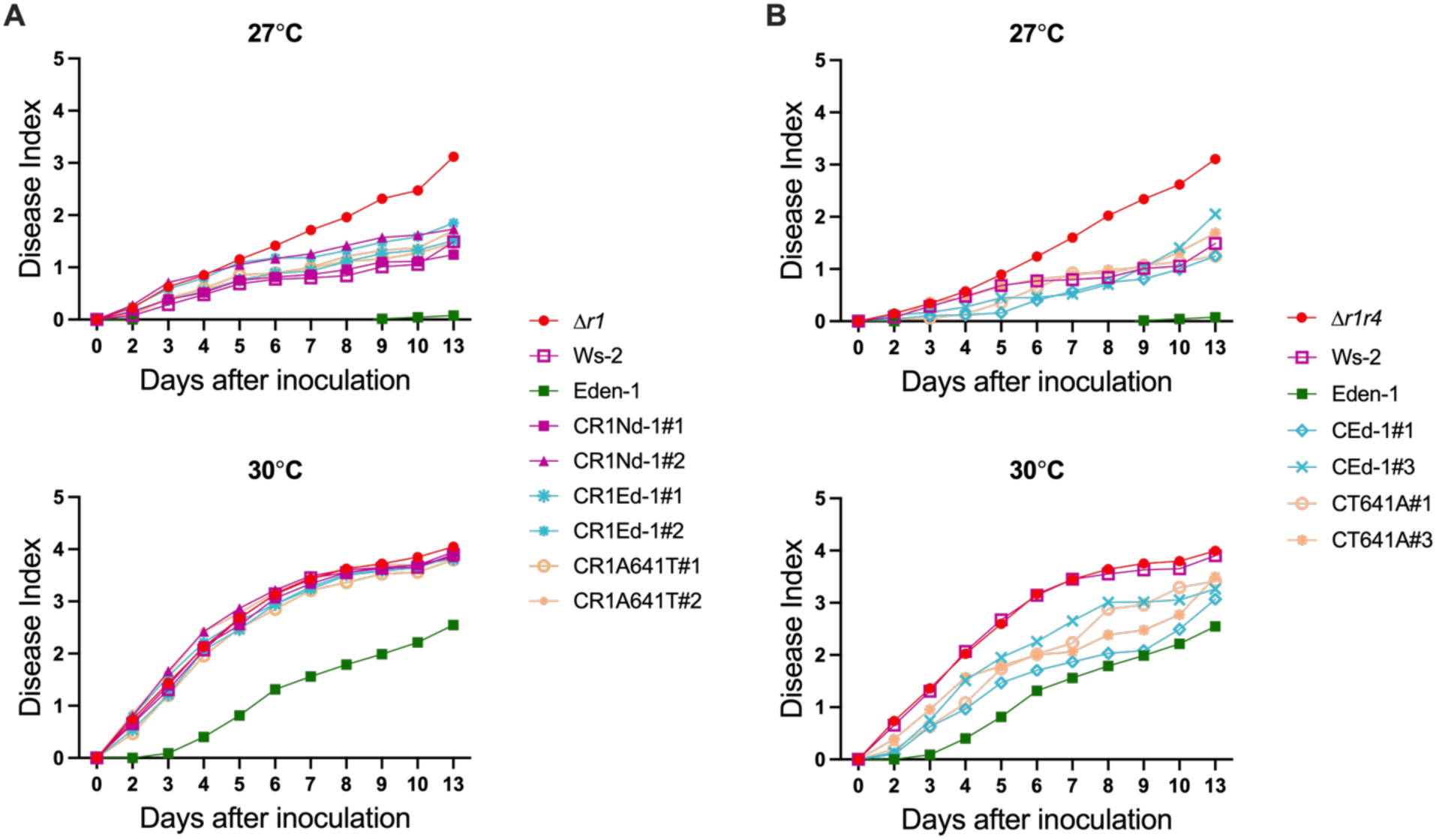
The introduction of the A641T substitution in *RRS1^Nd-1^* is not sufficient to fully restore thermostable immune response in *A. thaliana* to GMI1000 at 30°C. (**A**) The *rrs1-1* single mutant complemented with *RRS1^Nd-1^, RRS1^Eden-1^*, or *RRS1^Nd-1^ ^A641T^* genomic constructs, *rrs1-1* single mutant, and wild-type Eden-1 were tested for disease resistance to GMI1000 at 27°C and 30°C in two biological replicates. For each biological replicate, 18 to 20 plants were tested for each independent line. (**B**) The *rps4-21/rrs1-1* double knockout mutant complemented with *RPS4/RRS1^Eden-1^* haplotype (CEd-1#1, CEd-1#3), and *RPS4/RRS1^Eden-1^ ^T641A^* (with the reverse substitution T641A introduced into *RPS4/RRS1^Eden-1^;* CT641A#1, CT641A#2) were tested for disease resistance to GMI1000 at 27°C and 30°C in two biological replicates. For each biological replicate, 8 to 10 plants were tested for each independent line.

**Fig. S7.**
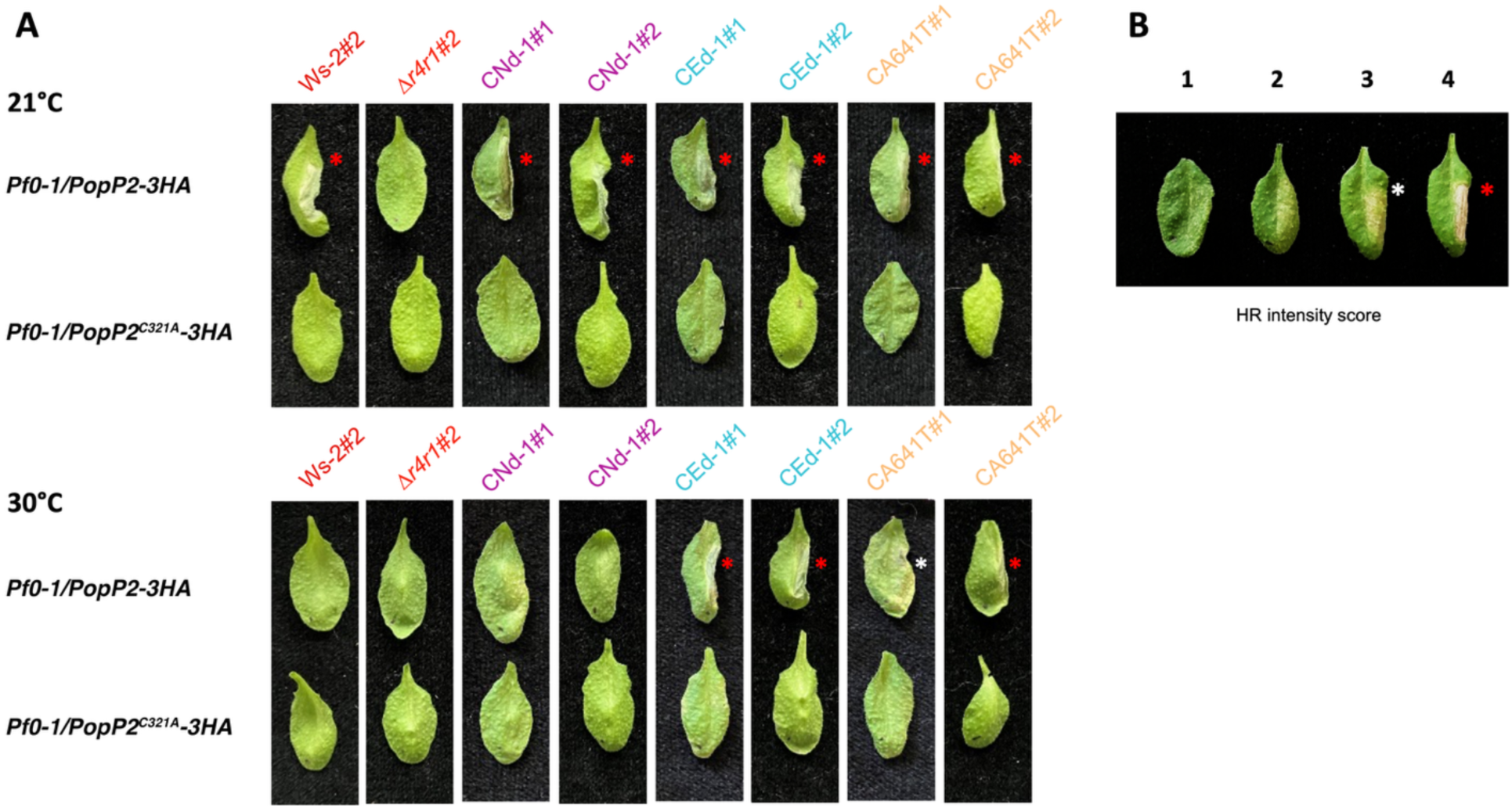
Temperature-dependent activation of RPS4/RRS1-mediated immune responses following *Pf0-1*-mediated delivery of PopP2. **(A)** Leaves from wild-type and genetic complement plants were infiltrated with *Pseudomonas fluorescens Pf0-1* strains delivering either wild-type PopP2-3HA or the catalytic mutant PopP2^C321A^-3HA, at 21°C and 30°C. Cell death responses were assessed 24 hours post-infiltration. Strong hypersensitive response (HR)-like cell death was observed in Ws-2, Eden-1, and all complemented lines (CEd-1#1, CEd-1#2, CNd-1#1, CNd-1#2, CA641T#1, CA641T#2) at 21°C upon PopP2 delivery, but not with the PopP2^C321A^-3HA mutant, confirming specific recognition by the RPS4/RRS1 complex. At 30°C, only lines complemented with *RPS4/RRS1^Eden-1^* and *RPS4/RRS1^Nd-1^ ^A641T^* retained the ability to trigger cell death in response to PopP2, indicating thermostable immune activation. The *rps4-21/rrs1-1* double mutant (*Δr4/r1*) and wild-type Ws-2 plants did not show a response under either condition at 30°C. (**B**) HR intensity was scored following a scale from 1 to 4. White and red asterisks indicate the presence of mild and strong HR, respectively.

**Fig. S8.**
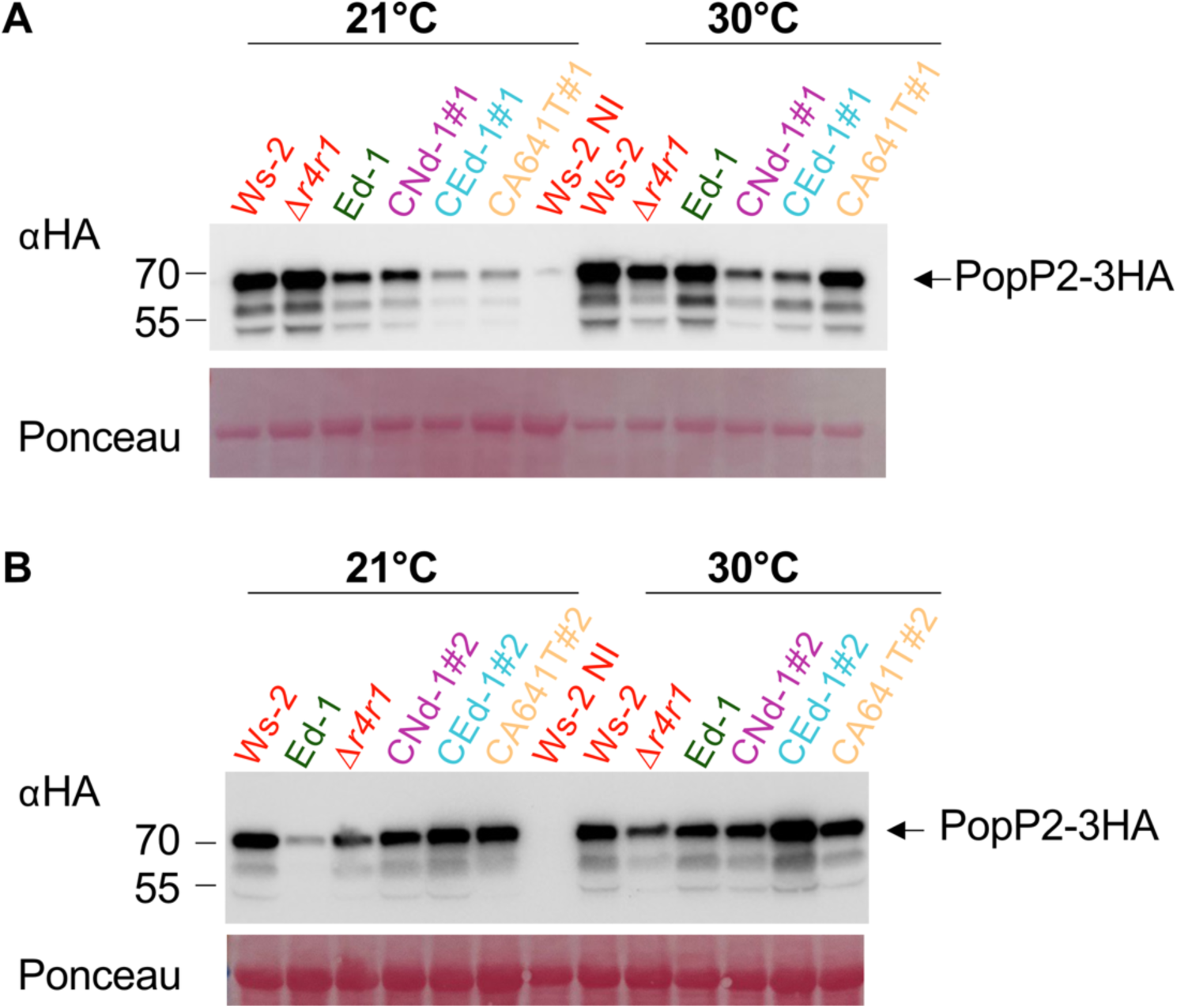
Immunodetection of PopP2-3HA and PopP2^C321A^-3HA delivered by *Pf0-1* in *A. thaliana* Ws-2, Eden-1, *Δr4/r1*, and transgenic lines at 21°C and 30°C. Total proteins were extracted 6 hours after leaf infiltration and analyzed by western blot using anti-HA antibodies. Ponceau Red staining was used as a loading control. *Δr4r1* refers to the *rps4-21/rrs1-1* double mutant and Ed-1 refers to Eden-1. NI refers to samples that were not infiltrated with *Pf0-1* cells.

**Fig. S9.**
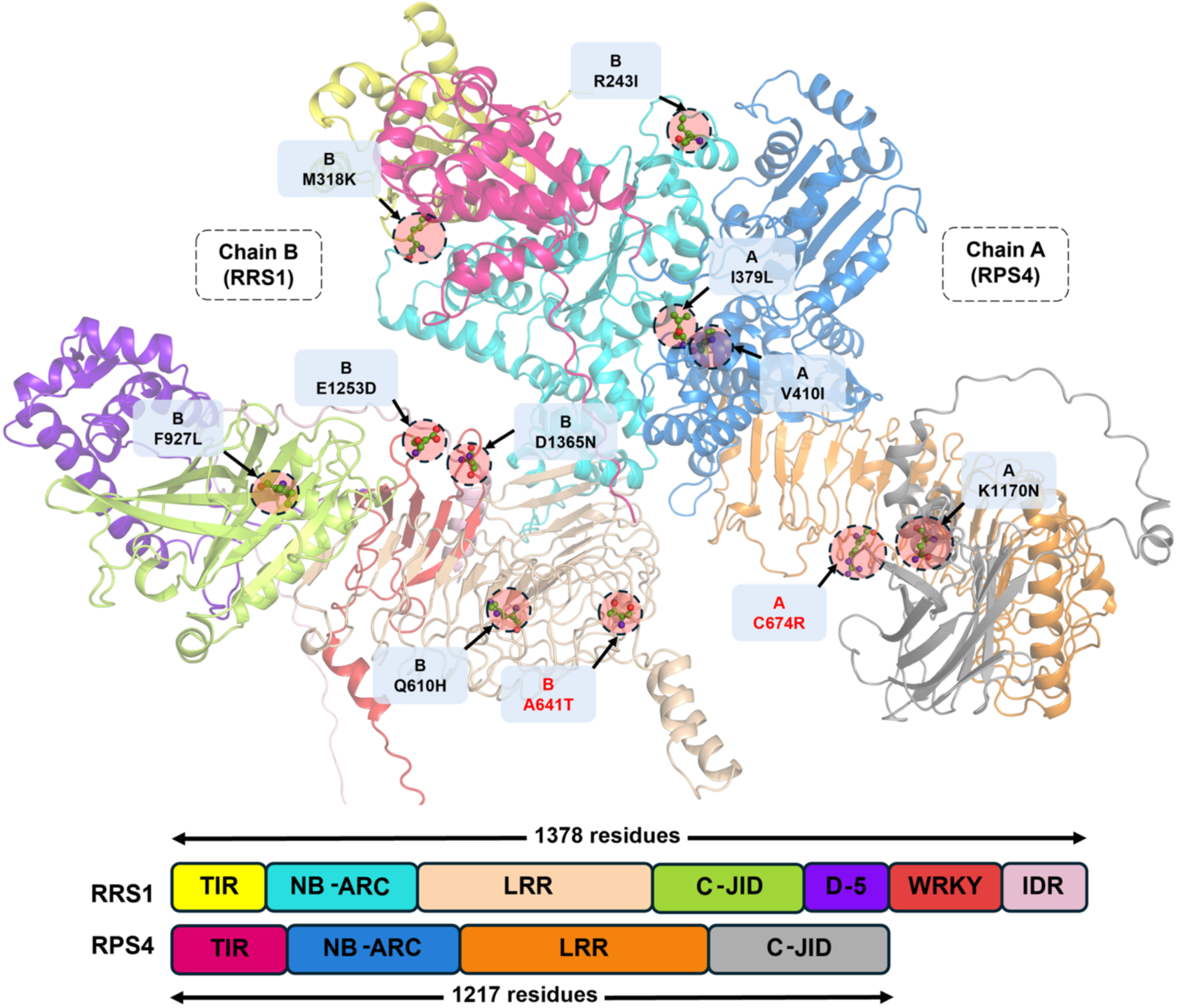
Eden-1 polymorphisms mapped onto the RPS4/RRS1^Nd-1^ structure. Amino acid polymorphisms specific to RPS4/RRS1^Eden-1^ were introduced into RPS4/RRS1^Nd-1^, whose structure was predicted using AlphaFold 3. RPS4 (chain A) and RRS1 (chain B) are shown in distinct colors, with domains colored as indicated in the schematic diagram. Polymorphic residues are highlighted as pink spheres, with non-synonymous polymorphisms labelled in red.

